# Quantifying Amide-Aromatic Interactions at Molecular and Atomic Levels: Experimentally-determined Enthalpic and Entropic Contributions to Interactions of Amide sp^2^O, N, C and sp^3^C Unified Atoms with Naphthalene sp^2^C Atoms in Water

**DOI:** 10.1101/2023.07.12.548600

**Authors:** Emily Zytkiewicz, Irina A. Shkel, Xian Cheng, Anuchit Rupanya, Kate McClure, Rezwana Karim, Sumin Yang, Felix Yang, M. Thomas Record

## Abstract

In addition to amide hydrogen bonds and the hydrophobic effect, interactions involving π-bonded sp^2^ atoms of amides, aromatics and other groups occur in protein self-assembly processes including folding, oligomerization and condensate formation. These interactions also occur in aqueous solutions of amide and aromatic compounds, where they can be quantified. Previous analysis of thermodynamic coefficients quantifying net-favorable interactions of amide compounds with other amides and aromatics revealed that interactions of amide sp^2^O with amide sp^2^N unified atoms (presumably C=O···H-N hydrogen bonds) and amide/aromatic sp^2^C (lone pair-π, n-π^*^) are particularly favorable. Sp^3^C-sp^3^C (hydrophobic), sp^3^C-sp^2^C (hydrophobic, CH-π), sp^2^C-sp^2^C (hydrophobic, π-π) and sp^3^C-sp^2^N interactions are favorable, sp^2^C-sp^2^N interactions are neutral, while sp^2^O-sp^2^O and sp^2^N-sp^2^N self-interactions and sp^2^O-sp^3^C interactions are unfavorable. Here, from determinations of favorable effects of fourteen amides on naphthalene solubility at 10, 25 and 45 °C, we dissect amide-aromatic interaction free energies into enthalpic and entropic contributions and find these vary systematically with amide composition. Analysis of these results yields enthalpic and entropic contributions to intrinsic strengths of interactions of amide sp^2^O, sp^2^N, sp^2^C and sp^3^C unified atoms with aromatic sp^2^C atoms. For each interaction, enthalpic and entropic contributions have the same sign and are much larger in magnitude than the interaction free energy itself. The amide sp^2^O-aromatic sp^2^C interaction is enthalpy-driven and entropically unfavorable, consistent with direct chemical interaction (e.g. lone pair-π) while amide sp^3^C- and sp^2^C-aromatic sp^2^C interactions are entropy-driven and enthalpically unfavorable, consistent with hydrophobic effects. These findings are relevant for interactions involving π-bonded sp^2^ atoms in protein processes.

**Table of Contents Graphic:** 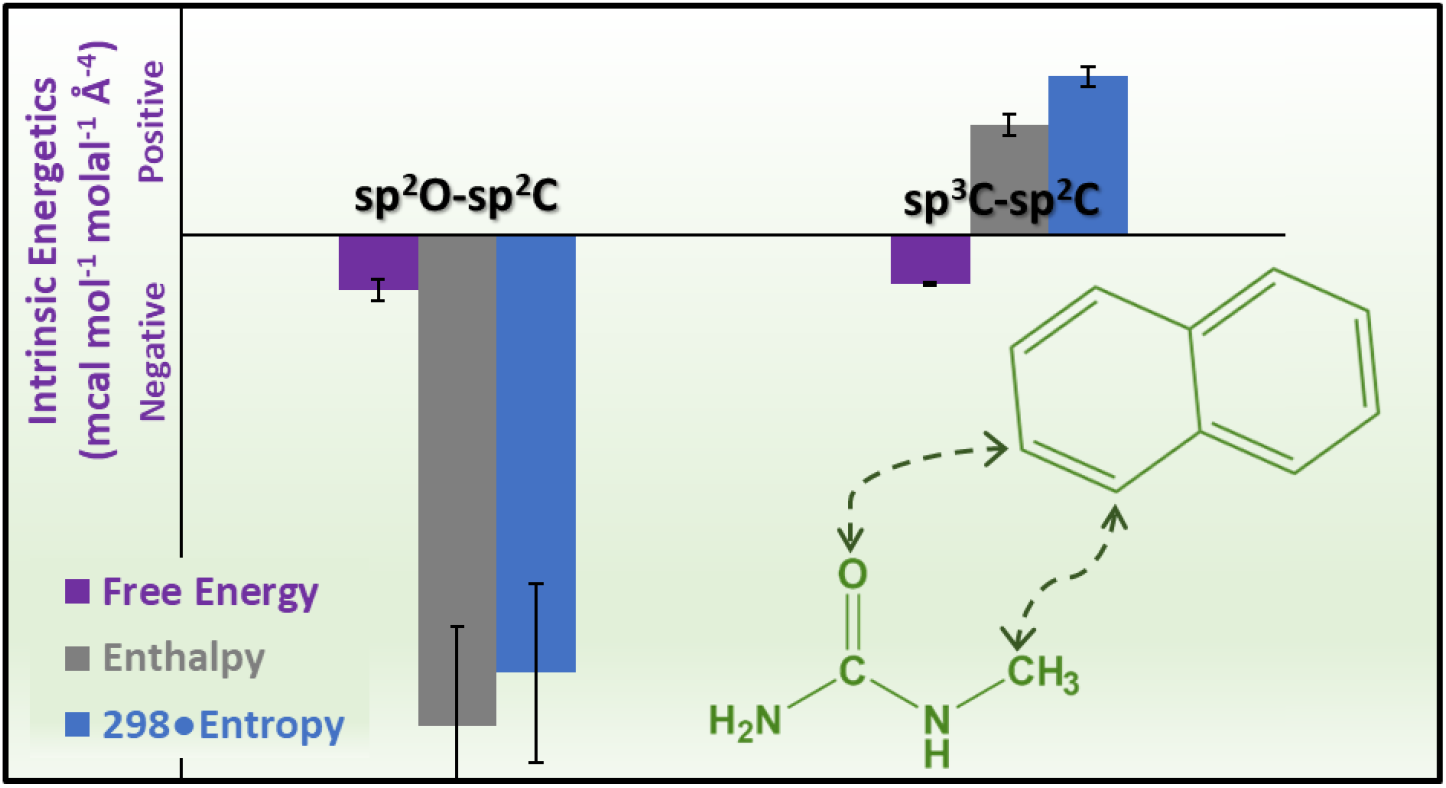

## Introduction

Self-assembly and binding interactions of biopolymers involve large numbers of relatively weak favorable (and unfavorable) interactions between individual atoms and/or groups. Because these individual interactions are comparable in strength to interactions of the atoms or groups with water, they have been difficult to quantify, resulting in ambiguity with regard to their contributions to the thermodynamics of forming biopolymer complexes and assemblies. These weak interactions also determine the effects of stabilizing or destabilizing biochemical solutes like glycine betaine or urea on biopolymer processes, and the strengths of interaction of these small solutes with one another in aqueous solution, where they are responsible for the deviations from a random solute mixture that are quantified as solute nonideality. From measurements of the effect of changing the concentration of one solute on the activity coefficient of a second solute, we determine information about the solute-solute interaction free energy. The quantity obtained is the chemical potential derivative (*∂*μ_2_/*∂*m_3_)_T,P,m2_ = μ_23_, a model-independent fundamental thermodynamic coefficient which can be interpreted as a transfer free energy and which quantifies the free energy of the preferential interaction of the two solutes, relative to their interactions with water.^1-3^

For interactions of two small solutes, solubility or osmometry assays are used to determine μ_23_.^3-14^ Solubility assays are used in the present work because one solute (naphthalene) is only sparingly soluble and because data as a function of temperature are needed. Where both solutes are sufficently soluble, osmometry is the assay of choice but only a few temperatures are readily available (e.g. 0 °C by freezing point osmometry; room temperature by vapor pressure osmometry). Analysis of these μ_23_ values demonstrates that interactions of individual groups or types of unified atoms on the surfaces of one or both of the solutes make additive contributions to μ_23_.^2,3,7-14^ Chemically-reasonable interaction parameters (called α-values) are obtained that quantify these contributions. An earlier approach determined the effect of urea and other solutes on the solubility of amino acids and dipeptides, and assumed additivity to obtain contributions of the peptide backbone and the nineteen amino acid side chains to a transfer free energy.^4-6^ The two approaches have been compared, and advantages of the approach used here have been discussed.^3^

Recent research determined preferential interaction free energies (μ_23_ values) for 105 amide-amide and amide-aromatic interactions^12,14^ and analyzed them as additive contributions from interactions of aliphatic sp^3^C and amide sp^2^O, N and C unified atoms of the amide compound with one another and with the sp^2^C of the aromatic compound (naphthalene, anthracene).^14^ From this analysis we obtained ten intrinsic interaction strengths (two-way α-values) for the contributions to μ_23_ from each pairwise interaction between these unified atoms, expressed per unit of water-accessible surface area (ASA). Intrinsic strengths of interactions of aromatic sp^2^C and amide sp^2^C with other atom types were found to be sufficiently similar that they could be analyzed together.^12,14^

Interactions of amide sp^2^O with amide sp^2^N unified atoms (sp^2^O-sp^2^N; presumably C=O···H-N hydrogen bonds^15-19^) and with amide/aromatic sp^2^C (sp^2^O-sp^2^C; presumably a lone pair-π^20-22^ or n-π^* 18,23-25^ interaction) are found to be particularly favorable.^14^ All pairwise C-C interactions investigated are favorable. These include interactions between sp^3^C unified atoms of two amide compounds (sp^3^C-sp^3^C), interactions between sp^3^C atoms of amide compounds and amide or aromatic sp^2^C unified atoms (sp^3^C-sp^2^C) and interactions of amide sp^2^C with aromatic sp^2^C (sp^2^C-sp^2^C).^14^ Sp^3^C-sp^3^C interactions are presumably hydrophobic.^26-30^ Sp^3^C-sp^2^C and sp^2^C-sp^2^C interactions may also involve the π system(s) (e.g. CH-π^31-35^ and π-π^36-38^ interactions).

Interactions between amide sp^2^N and sp^3^C unified atoms of amide compounds (sp^2^N-sp^3^C) are modestly favorable while amide sp^2^N-sp^2^N self-interactions are unfavorable,^14^ consistent with the observation that amide sp^2^N-sp^2^N hydrogen bonds are uncommon in proteins.^39^ Self-interactions between amide sp^2^O unified atoms (sp^2^O-sp^2^O) are particularly unfavorable, as are sp^2^O-sp^3^C interactions.^14^ These unfavorable interactions of amide sp^2^O unified atoms with other amide sp^2^O unified atoms and with sp^3^C unified atoms of amide compounds indicate the strong preference of amide sp^2^O to interact with water than with itself or sp^3^C. Likewise amide sp^2^N unified atoms evidently prefer to interact with water than with themselves.

From these α-values and ASA information, one can predict the interaction of any amide solute with another amide or aromatic solute. Because the protein surface exposed in unfolding is composed mostly of these hydrocarbon and amide unified atoms, effects of any amide solute on the stability of a protein to unfolding can also be predicted from information about the amount and composition of the ΔASA of unfolding.^14^

Here solubility assays at 10, 25 and 45 °C are used to determine effects of temperature on free energies of preferential interactions (μ_23_ values) of fourteen amide compounds with naphthalene. Global analysis of these data for each amide compound yields s_23_, the entropic contribution to μ_23_, which is the negative of the derivative of μ_23_ with respect to temperature.

From μ_23_ and s_23_ we obtain enthalpic (h_23_) contributions to each amide-naphthalene preferential interaction (h_23_ = μ_23_ + Ts_23_). We interpret these results to obtain the four two-way α-values and their entropic and enthalpic components for interactions of the sp^2^O, sp^2^N, sp^2^C and sp^3^C unified atoms of amide compounds with aromatic sp^2^C atoms. These α-values make good chemical sense and, moreover, are useful to predict the μ_23_ value and its entropic (s_23_) and enthalpic (h_23_) components for any amide-aromatic interaction.

To examine whether an atom-atom (four-parameter) analysis of the free energy, enthalpy and entropy of amide-naphthalene interactions is more justifiable than alternatives, we also interpret these data using two interaction parameters, one quantifying amide π-aromatic π interactions, the other for amide sp^3^C-aromatic π (i.e. CH–π) interactions. We compare the statistical and chemical significance of free energy α-values and their enthalpic and entropic components obtained using these two analyses of amide-naphthalene interactions, and conclude that the atom-atom (four-parameter) analysis is statistically better and yields interaction parameters (two-way α-values) that are more chemically meaningful.

Combining the 25 °C amide-naphthalene data obtained here with that reported previously for amide-amide and amide-aromatic interactions,^14^ we analyze this large (109 member) data set to refine the ten two-way α-values obtained previously that quantify the interactions of amide sp^2^O, amide sp^2^N, aliphatic sp^3^C and amide/aromatic sp^2^C atoms. Additionally, as a test of whether an atom-atom (ten-parameter) analysis of the free energy of amide-amide and amide-aromatic interactions is more justifiable than alternatives, we also analyze the 109-member set of amide-amide and amide-aromatic μ_23_ values as π-π and CH-π interactions. This analyses requires only four two-way α-values, less than half the number needed for an atom-atom analysis (ten two-way α-values). Although π-π and CH-π α-values obtained from this fit provide moderately good predictions of μ_23_ values from ASA information, neither the statistics of the fit nor the chemical significance of the parameters is of the same quality as when interpreted using ten atom-atom interactions.

### Background on Determinations of Molecular and Atom-Atom Amide-Naphthalene Interaction Free Energies; Extension to Interaction Entropies and Enthalpies

#### Solubility Determinations of Naphthalene Transfer Free Energies from Water to Amide Solutions 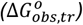 and their Entropic 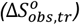 and Enthalpic 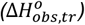 Components

Early research investigated interactions of amino acids (solute species 2) with urea or osmolytes (solute species 3) by solubility measurements. In these studies the effect of solute 3 on the solubility of solute 2 was determined and used to calculate an observed standard free energy of transfer 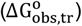 of solute 2 from water to a specified concentration of solute 3:^4-6^

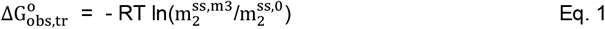

where 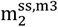 is the molal scale solubility of solute 2 at molality m_3_ of solute 3 and 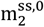 is the solubility in absence of solute 3. In the studies reported here naphthalene is solute 2 and a set of 14 amides, including urea, are investigated as solute 3. Values of 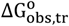 determined as a function of amide concentration m_3_ provide information about the change in the activity coefficient of naphthalene as a function of m_3_, and therefore about the amide-naphthalene preferential interaction free energy μ_23_ (see SI Eqs. S1-3).

The entropy of transfer 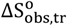 is obtained from the temperature dependence of 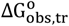

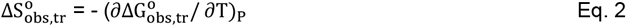

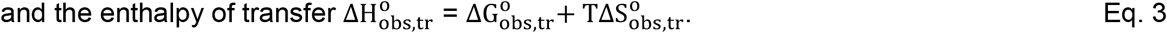

No systematic studies of 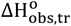 or 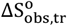 have been reported previously.

#### Free Energies of Amide-Naphthalene Preferential Interaction (μ_23_) and their Entropic (s_23_) and Enthalpic (h_23_) Components, Determined from Transfer Free Energies

The observed transfer free energy 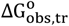 depends on the concentration of solute 3. Expanding 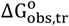 in a Taylor-Maclaurin series in powers of m_3_ yields:

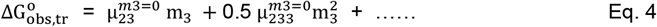

In Eq. 4, µ_23_ and µ_233_ are defined as

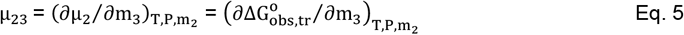

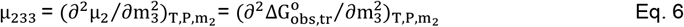

(see SI Eqs. S2-3), and 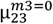 and 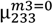 are limiting values of µ_23_ and µ_233_ as m_3_ approaches 0.

The chemical potential derivative μ_23_ quantifies the preferential interaction of the amide compound with naphthalene. Since μ_23_ is the initial slope of a plot of 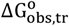 vs. m_3_, it is the solubility *m*-value. Since all μ_23_ and μ_233_ obtained here are 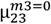 and 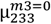 we omit this superscript hereafter. Values of μ_23_ at temperature T are related to those at 298 K by

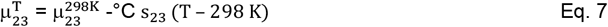

where s_23_ = - (∂μ_23_/ ∂T)_P_ is the entropy of the naphthalene-amide preferential interaction, assumed independent of T. Since μ_23_ refers to 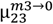, similarly s_23_ is the entropy of the preferential interaction in the limit as m_3_ → 0.

From Eqs. 4 and 7,

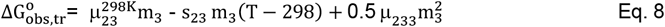

Naphthalene solubility data as a function of concentration of each amide compound at 10, 25 and 45 °C are fit globally to Eq. 8 to obtain 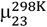, s_23_ and μ_233_, where μ_233_ is also assumed to be independent of T. The enthalpy of preferential interaction (also assumed independent of T) is obtained from 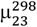 and s_23_.

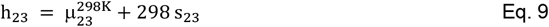

Alternatively, data for 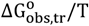 as a function of m_3_ and T may be analyzed globally using Eq. S4 (paralleling Eq. 8) to obtain h_23_ directly. Values of h_23_, s_23_ and 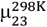 obtained from Eq. S4 and Eq. 9 are the same within uncertainty as those obtained from Eqs. 8-9.

#### Amide Atom-Naphthalene Atom Contributions (α_ij_-values) to Preferential Interaction Free Energies; Proposed Extension to Preferential Interaction Enthalpies and Entropies

Values of μ_23_ for amide-naphthalene interactions are dissected into interactions of naphthalene and its sp^2^C atoms with the four types of amide atoms (amide sp^2^O, sp^2^N and sp^2^C; methyl or methylene sp^3^C), listed by the index i (1 ≤ i ≤ 4), using the relationship^14^

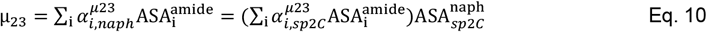

where 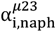 and 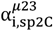 are the intrinsic strengths of interaction of amide atom type i with naphthalene and with its sp^2^C atoms, both expressed per unit of water-accessible surface area (ASA) of amide atom i (cf. Table S1 for ASA values). In Eq. 10, 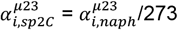 where the 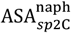 is 273 Å^2^. The assumptions of additivity in the one-way and two-way analyses of Eq. 10 have been tested and justified previously^3,7-14^ and are also tested here, because the size of the experimental data set (14 μ_23_ values) exceeds the number of fitting parameters (four different 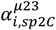).

Here (in Discussion) we apply the two way analysis of Eq. 10 to amide-aromatic intereactions and extend this analysis to h_23_ and s_23_. We introduce and test dissections of s_23_ and h_23_ based on additivity and ASA that are analogous to Eq. 10 for 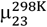. From this analysis we obtain enthalpic 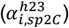 and entropic 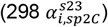 contributions to free energy α-values 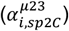 from interactions of naphthalene sp^2^C unified atoms with amide sp^2^O, sp^2^N and sp^2^C and methyl or methylene sp^3^C atoms of the amide compounds. We also use a similar approach to examine whether interactions with the amide group can be successfully modeled as interactions with the amide π system.

## Results

### Large Effect of Increasing Hydrocarbon (sp^3^C) ASA of Amide Compounds on the Temperature-Dependence of the Free Energy 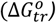 of Transfer of Naphthalene from Water to Amide Solutions

To quantify the thermodynamics of amide-naphthalene interactions, naphthlene solubilities were measured as a function of molal concentration m_3_ of fourteen different amide compounds at three temperatures (10, 25, 45 °C). Naphthalene transfer free energies 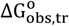 (Eq. 1) calculated from these solubilities are plotted vs m_3_ in Figures 1 and S1.

**Figure 1.**
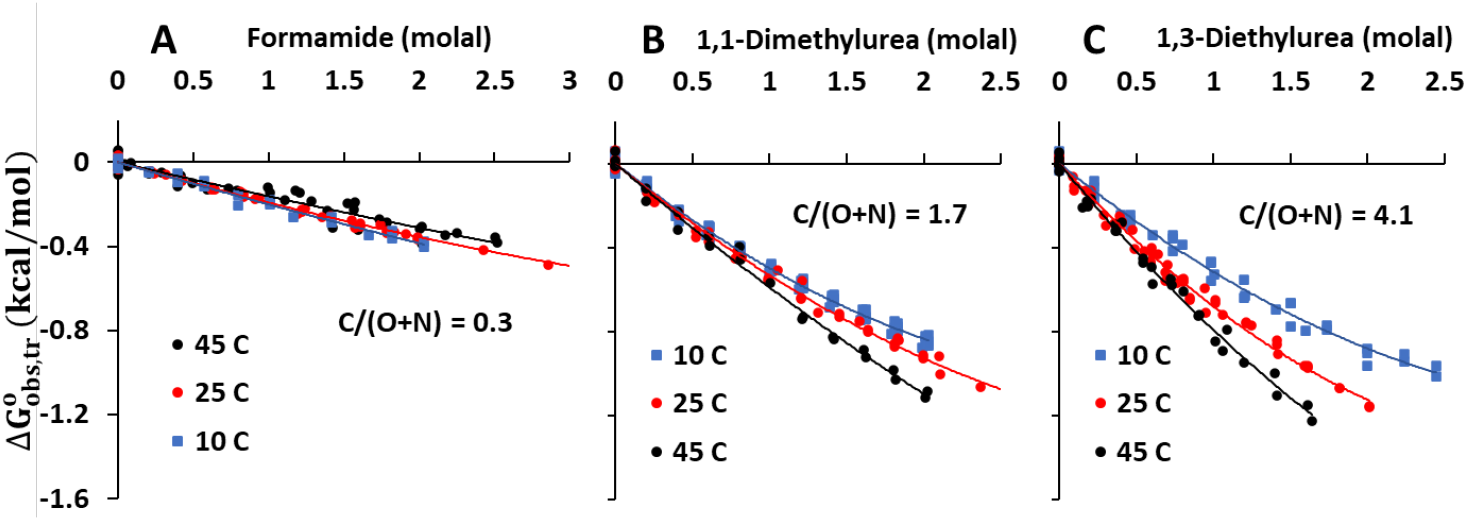
Naphthalene transfer free energies 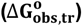 from water to amide solutions at 10, 25, and 45 °C: different temperature effects for different amide nonpolar (C) : polar (O, N) ASA ratios. Values of 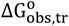, obtained from solubilities (Eq. 1) at 10, 25, and 45 °C are plotted vs. molal concentration of three amides (panel A formamide, panel B 1,1-dimethylurea, and panel C 1,3-diethylurea) selected for their widely different ratios of nonpolar (C) to polar (O+N) surface area, indicated as C/(O+N) in the figure panels. Initial slopes of quadratic fits of the data of each panel to Eq. 4 (plotted curves) are μ23 values at 10, 25, and 45 °C (see Table 1). Analogous plots for 11 other amide compounds are given in Figure S1.

Results for three amide compounds (formamide, 1,1-dimethylurea (1,1-dmu) and 1,3-diethylurea (1,3-deu)), selected for their very different amounts of hydrocarbon (methyl and methylene sp^3^C) and amide (sp^2^O, sp^2^N, sp^2^C) surface area (ASA) and their different ratios of nonpolar (C) to polar (O+N) ASA, are shown in Figure 1. These are plotted using the same free energy and concentration scales so the magnitudes and temperature dependences of 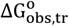 for the three amides can be visually compared. Corresponding plots for eleven other amide compounds, together with an expanded free energy scale plot for formamide, are given in the panels of Figure S1.

**Table 1.**
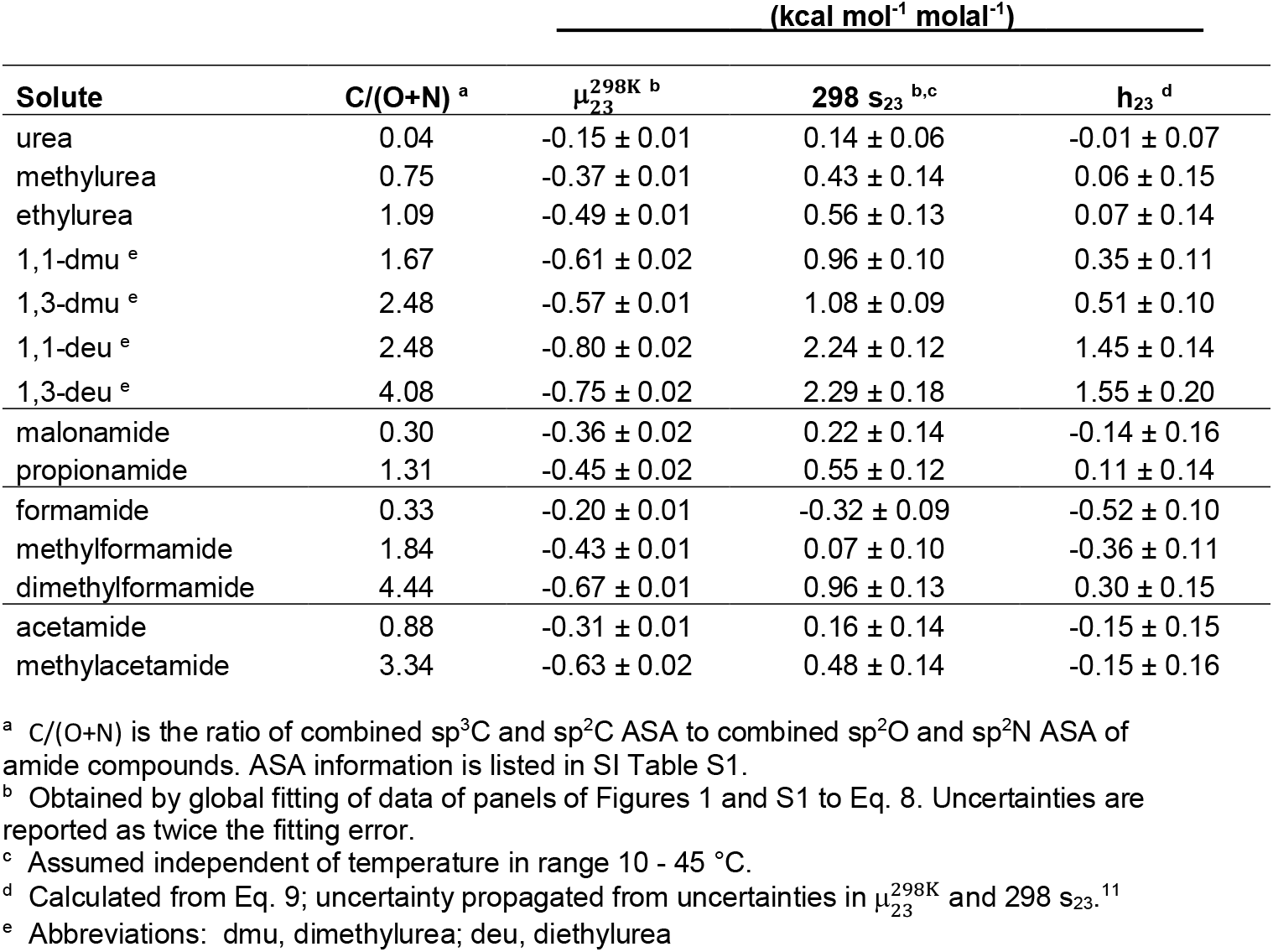
Thermodynamics of Amide-Naphthalene Preferential Interactions.

All fourteen amide compounds increase the solubility of naphthalene at all three temperatures, indicating that all of them interact favorably with naphthalene. Values of 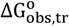 for transfer of naphthalene from water to these amide solutions are negative and increase with increasing amide concentration. All plots in Figures 1 and S1 are linear within uncertainty at amide concentrations below 0.5 molal. From Eq. 4-5 the initial slopes of these plots are free energies of amide-naphthalene preferential interactions (μ_23_ values). Except for the smallest amides (formamide, urea), curvature is observed at amide concentrations above 0.5 -1.0 molal. This curvature, quantified by the derivative of μ_23_ with respect to m_3_ (μ_233_; Eq. 6) generally increases with the size of the amide compound.

Formamide is the smallest amide compound, lacks alkyl (sp^3^C) hydrocarbon, and has a relatively small C/(O+N) ASA ratio of 0.3 (Table 1). Figure 1 shows that the transfer free energy 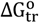 of naphthalene from water to a specified formamide concentration is much smaller in magnitude (less favorable) and much less temperature-dependent than for the other amides in Figure 1. From Figure 1A (expanded in Figure S1H) the transfer free energy 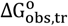 of naphthalene from water to a specified formamide concentration is slightly more favorable at 10 °C than at 25 °C, and least favorable at 45 °C. Because 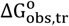 increases with increasing temperature, the corresponding entropy of transfer 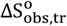 of naphthalene from water to formamide is negative (unfavorable) and the favorable 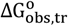 therefore results entirely from a favorable (negative) enthalpy of transfer 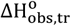.

Formamide is one of five amide compounds in Figure S1 for which naphthalene transfer free energies are only weakly temperature dependent. Of the other four (urea, acetamide, malonamide, methylformamide), all but methylformamide also have relatively small C/(O+N) ASA ratios (Table 1). Unlike formamide, 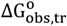 for each of these four amides, compared at any amide concentration, decreases (becomes slightly more negative) with increasing temperature, revealing that the corresponding entropies of transfer 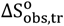 of naphthalene from water to these amides are positive (favorable). Values of 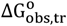 for these four amides are 1.5-to 2-fold larger in magnitude than those of formamide at any amide concentration.

Alkylated ureas with sp^3^C (methyl, methylene) ASA have much larger C/(O+N) ASA ratios than formamide and, compared at the same concentration, are much better solubilizing agents for naphthalene than formamide. Two examples are shown in Figure 1B-C. Values for 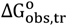 for transfer of naphthalene from water to 1,1-dmu (C/(O+N) = 1.7) and to 1,3-deu (C/(O+N) = 4.1) are three to four times more favorable than for formamide or urea at the same temperature and m_3_. For these and other alkylureas in Figure S1, 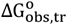 decreases strongly with increasing temperature at any fixed m_3_, and the effect of temperature is larger for the larger alkylureas. Hence values of 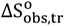 are positive and increase with increases in the C/(O+N) ASA ratio for these alkylureas.

Taken together, the results of Figures 1 and S1 indicate that the favorable 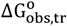 of naphthalene from water to amide solutions is the result of favorable interactions of naphthalene with both the polar (O+N) and nonpolar (C) portions of these amides. The favorable contribution from the nonpolar (C) portion of the amide compound to naphthalene 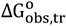 results from a very favorable contribution to 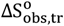, partially countered by an unfavorable 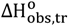. On the other hand, the origin of the favorable contribution from the polar (O+N) portion of the amide to naphthalene 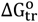 results from a favorable contribution to 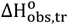.

### Quantifying Free Energies of Amide Compound-Naphthalene Preferential Interactions (μ_23_) and their Entropic and Enthalpic Parts (s_23_, h_23_) from 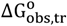

Free energies 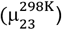 and entropies (s_23_) of the preferential interaction of each of fourteen amide compounds with naphthalene interactions are obtained from global fitting of naphthalene 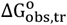 data (Figure 1 and S1) at 10, 25 and 45 °C to Eq. 8. Values of 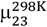 and 298 s_23_ are listed in Table 1, together with C/(O+N) ASA ratios for these amide compounds. Enthalpies (h_23_) of the preferential interactions of each amide compound with naphthalene, calculated from 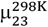 and 298 s_23_ by Eq. 9, are also listed in Table 1.

Alternatively, values of 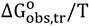 are analyzed as a function of 1/T using Eq. S4 to obtain 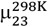 and h23, from which s23 is calculated by Eq. 9. Identical values of 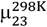 for interactions of each of the 14 amides with naphthalene are obtained by fitting to either Eq. 8 or S4. Similar but not identical values and uncertainties are obtained for h_23_ and s_23_ by these two approaches; these are compared in SI Tables S2 (for 298 s_23_) and S3 (for h_23_). Values of μ_233_ obtained from fitting the 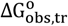 values to Eq. 8 or S4 are compared in Table S4. No significant differences in μ_233_ from these two fitting approaches are observed. For the alkylurea series, μ_233_ increases from approximately zero for urea to 0.19 for 1,3-diethylurea. Figure 2 and Table 1 group the fourteen amides into ureas, formamides, acetamides, and other amides, and list the compounds in each group in order of increasing C/(O+N) ASA ratio. Values of 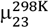, h_23_ and 298 s_23_ for interactions of each amide with naphthalene are plotted on the bar graph in Figure 2, which shows three patterns in the thermodynamics of these amide-naphthalene preferential interactions.

**Figure 2.**
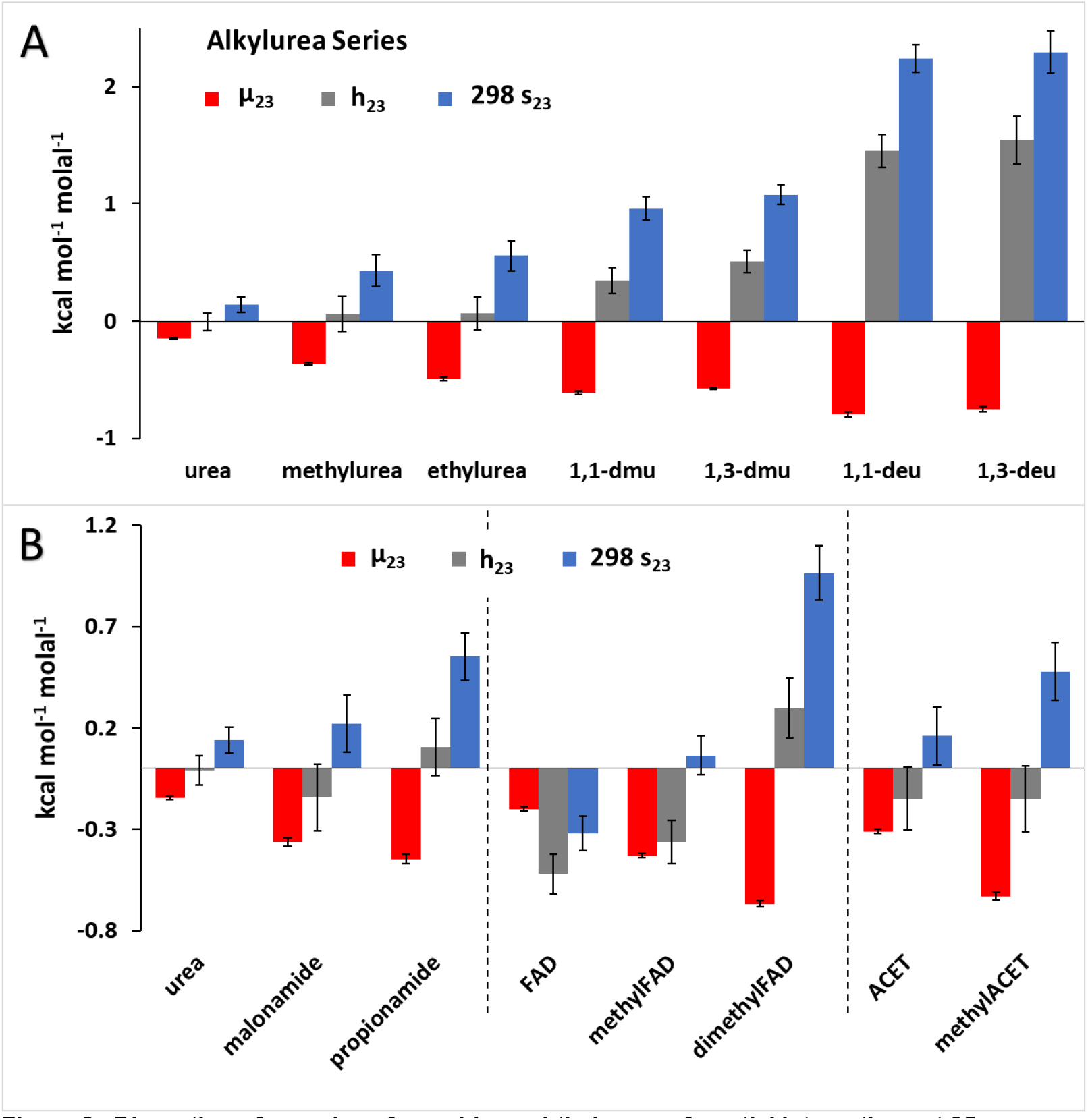
Dissection of μ_23_ values for amide-naphthalene preferential interactions at 25 °C into enthalpic (h_23_) and entropic (298 s_23_) contributions. Values of 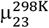, h_23_ and 298 s_23_ are listed in Table 1). Panel A shows the series of alkylureas in order of increasing C/(O+N) ASA ratio. Panel B shows three different groups of amides, also ordered within each group by increasing C/(O+N) ASA ratio. Abbreviations: FAD, formamide; ACET, acetamide.

i. Free energies 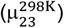 of these interactions, always favorable 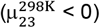, generally become more favorable with increasing C/(O+N) ASA ratio in each group. Minor exceptions are observed for the 1,3-dialkylureas, for which 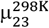 values are somewhat less favorable than for the corresponding 1,1-dialkylureas, even though the C/(O+N) ASA ratios are larger for 1,3-than for 1,1-dialkylureas. Within each group of amides, the changes in the C/(O+N) ASA ratio directly reflect an increase in aliphatic sp^3^C ASA and a reduction in amide sp^2^N ASA, with smaller reductions in the ASA of amide sp^2^O and sp^2^C (Table S1). The trends in μ_23_ in all series are therefore consistent with our previous findings that interaction of amide sp^3^C with aromatic sp^2^C is favorable while interaction of amide sp^2^N with aromatic sp^2^C is marginally unfavorable.^14^
ii. For the series of ureas and formamides and the two acetamides in Figure 2, enthalpic (h_23_) and entropic (298 s_23_) contributions to the amide-naphthalene preferential interaction free energy 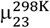 increase with increasing C/(O+N) ASA ratio. Enthalpies h_23_ are zero or negative (favorable) for the member of each group with the smallest C/(O+N) ASA ratio (urea, formamide, methylacetamide) and increase monotonically with increasing C/(O+N) ASA ratio. These interaction enthalpies are unfavorable for all alkylureas and for two other amides with moderately large C/(O+N) ratios (propionamide, dimethylformamide). Figure 2 reveals several general trends characteristic of these groups of amides. Entropic contributions to 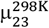 (i.e. 298 s_23_) are favorable (s_23_ > 0) for all amides except formamide, and increases monotonically with increasing C/(O+N) ASA ratio. These positive 298 s_23_ and the large increases in 298 s_23_ with increasing C/(O+N) ASA ratio are consistent with a hydrophobic origin of the favorable interaction of amide sp^3^C with aromatic sp^2^C because (at least for the liquid hydrocarbon model) the hydrophobic effect is predominantly entropic near room temperature.^26-28^
iii. The favorable preferential interactions of all alkylureas with naphthalene at 25 °C 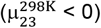 are entropy-driven (298 s_23_ > h_23_ > 0). (This situation also holds for alkylurea-naphthalene interactions at 10 and 45 °C.) Favorable preferential interactions between naphthalene and the most nonpolar amides in other groups in Figure 2 (dimethylformamide, propionamide, methylacetamide) are also entropy-driven with unfavorable (or modestly favorable) enthalpies. Favorable preferential interactions between naphthalene and the most polar amides in these groups (formamide, methylformamide, acetamide) are entirely or primarily enthalpy-driven, with an entropic contribution that is unfavorable for formamide and slightly favorable for methylformamide and acetamide.

## Analysis and Discussion

### Atom-atom Interpretation of Amide-Naphthalene Interaction Free Energies

As in a previous analysis of amide interactions,^14^ we use Eq. 10 to interpret the fourteen amide-naphthalene interaction free energies at 25 °C (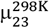; Table 1) as sums of contributions from interactions of the sp^2^O, sp^2^N, sp^2^C and sp^3^C unified atoms displayed on each amide compound with naphthalene sp^2^C unified atoms. In this analysis, Eq. 10 is written for each individual amide-naphthalene 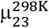, incorporating the appropriate ASA of sp^2^O, sp^2^N, sp^2^C and sp^3^C unified atoms of an amide compound (Table S1), the ASA of naphthalene sp^2^C (273 Å^2^; Table S1) and the four unknown amide atom-napthalene atom interaction strengths 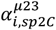. These 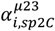 are contributions to 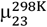 from interactions of 1 Å^2^ of amide compound atom-type I (where the index i represents sp^2^O, sp^2^N, sp^2^C, or sp^3^C unified atoms) with 1 Å^2^ of naphthalene sp^2^C ASA. Results are shown in Figure 3 and in the first column of Table 2. Very similar results are obtained from analysis of the combined set of 24 amide-aromatic (naphthalene, anthracene) 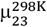 values using Eq. 10 (see Table S9 column 1).

**Table 2.**
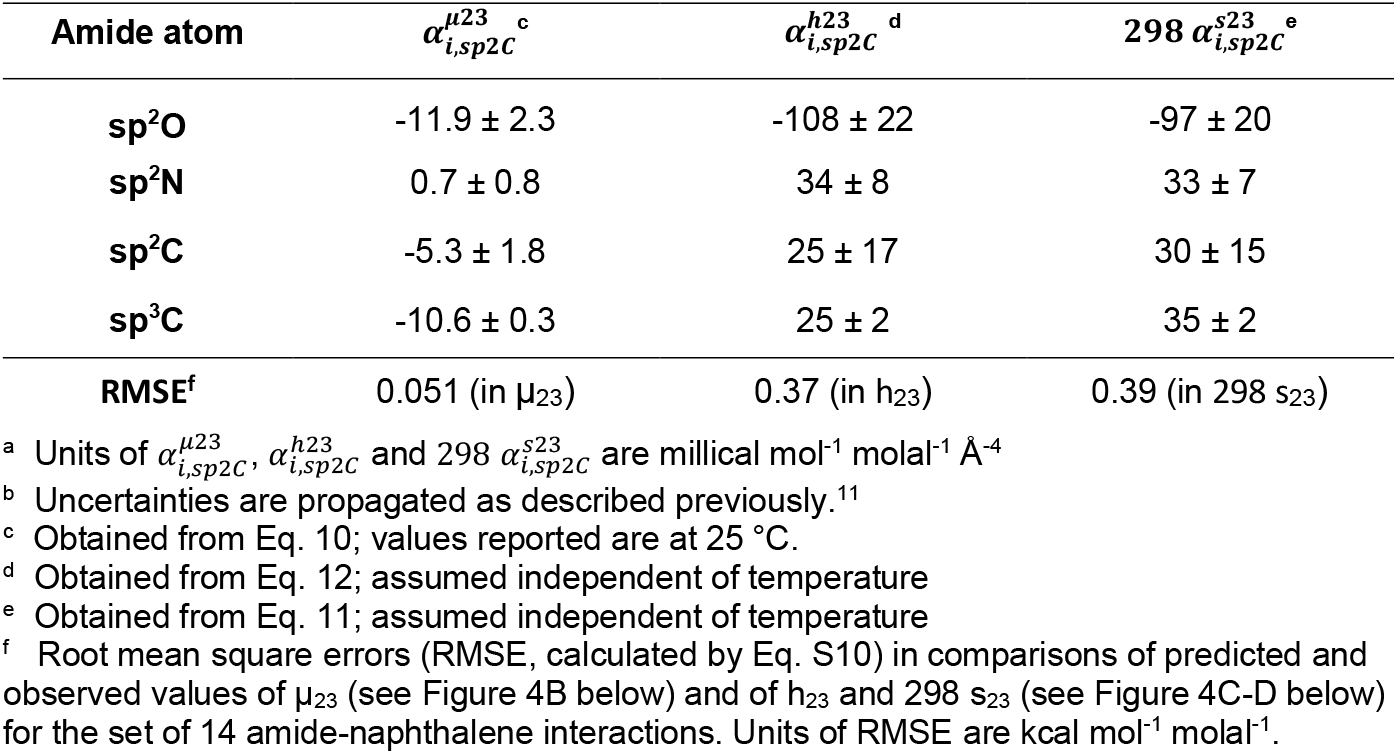
Thermodynamics of amide atom–naphthalene sp^2^C interactions: free energy α-values 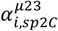 and their enthalpic 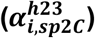 and entropic 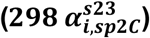 components ^a,b^.

| Amide atom | $\alpha_{i,sp2C}^{\mu23}$<sup>c</sup> | $\alpha_{i,sp2C}^{h23}$<sup>d</sup> | 298 $\alpha_{i,sp2C}^{s23}$<sup>e</sup> |
| --- | --- | --- | --- |
| sp<sup>2</sup>O | -11.9 ± 2.3 | -108 ± 22 | -97 ± 20 |
| sp<sup>2</sup>N | 0.7 ± 0.8 | 34 ± 8 | 33 ± 7 |
| sp<sup>2</sup>C | -5.3 ± 1.8 | 25 ± 17 | 30 ± 15 |
| sp<sup>3</sup>C | -10.6 ± 0.3 | 25 ± 2 | 35 ± 2 |
| RMSE<sup>f</sup> | 0.051 (in $\mu_{23}$) | 0.37 (in $h_{23}$) | 0.39 (in 298 $s_{23}$) |

<sup>a</sup> Units of $\alpha_{i,sp2C}^{\mu23}$ , $\alpha_{i,sp2C}^{h23}$ and 298 $\alpha_{i,sp2C}^{s23}$ are millical mol<sup>-1</sup> molal<sup>-1</sup> Å<sup>-4</sup>
<sup>b</sup> Uncertainties are propagated as described previously.<sup>11</sup>
<sup>c</sup> Obtained from Eq. 10; values reported are at 25 °C.
<sup>d</sup> Obtained from Eq. 12; assumed independent of temperature
<sup>e</sup> Obtained from Eq. 11; assumed independent of temperature
<sup>f</sup> Root mean square errors (RMSE, calculated by Eq. S10) in comparisons of predicted and observed values of $\mu_{23}$ (see Figure 4B below) and of $h_{23}$ and 298 $s_{23}$ (see Figure 4C-D below) for the set of 14 amide-naphthalene interactions. Units of RMSE are kcal mol<sup>-1</sup> molal<sup>-1</sup>.

**Figure 3.**
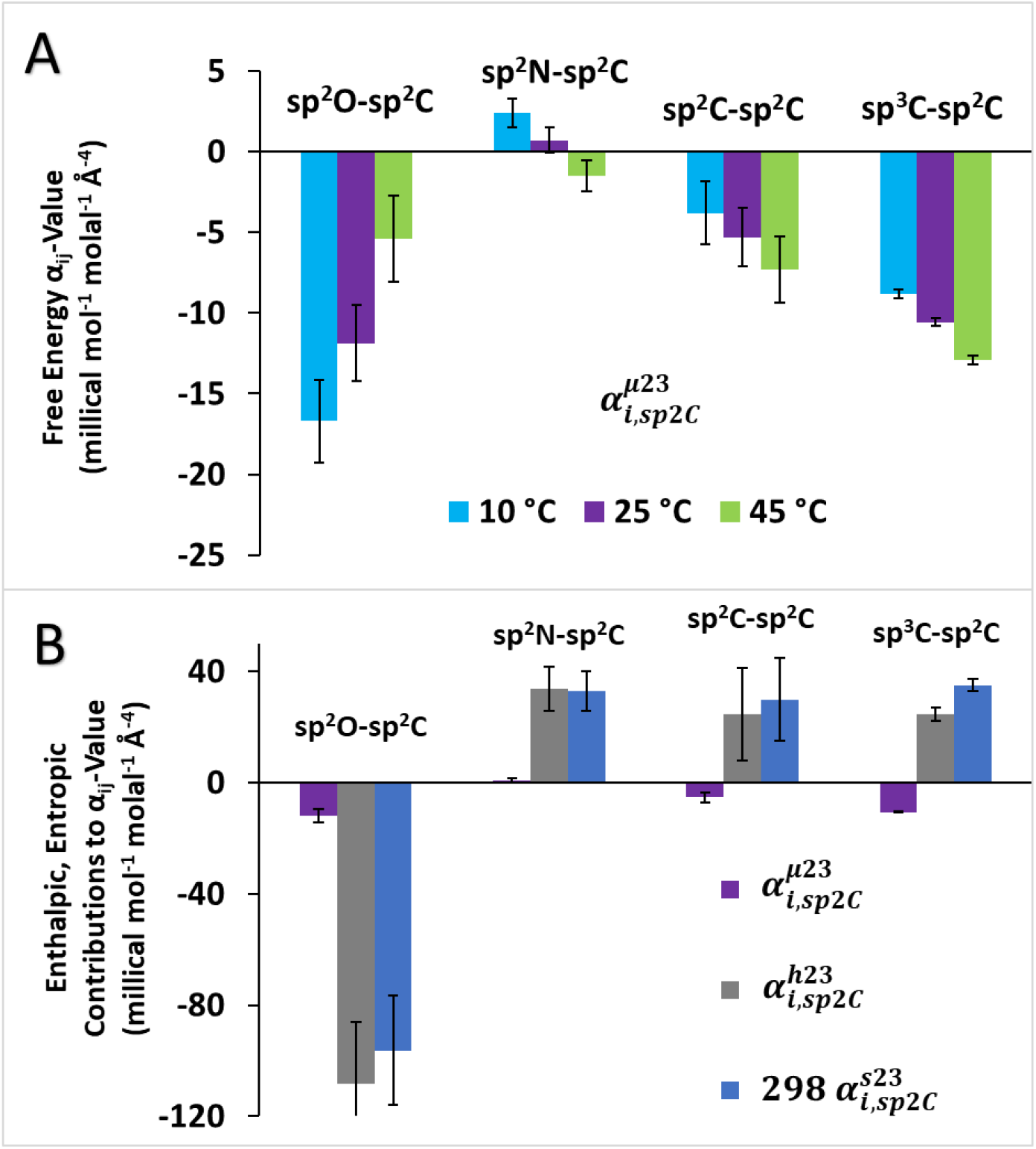
Two-way α-values for amide atom-naphthalene atom interactions: Temperature dependence (panel A) and enthalpic-entropic dissection (panel B). Panel A displays the four two-way α-values 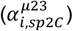 for amide atom-naphthalene sp^2^C interactions at 10, 25, and 45 °C. Panel B gives the dissections of these 25 °C 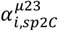 into enthalpic 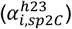 and entropic 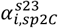contributions. All 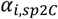 values and uncertainties are from Table 2.

This α-value analysis reveals that amide-naphthalene interactions are favorable (µ_23_ < 0; Figure 2) in the temperature range investigated because interactions of three of the four atom types of amide compounds with naphthalene sp^2^C are favorable at these temperatures. Intrinsic strengths 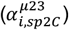 of interactions of amide sp^2^O and methyl/methylene sp^3^C with naphthalene sp^2^C are both very favorable, ranking with the amide sp^2^N-amide sp^2^O interaction (the amide NH···O=C hydrogen bond^14^) as the intrinsically most favorable amide interactions investigated. We previously proposed that the very favorable interaction of amide sp^2^O with aromatic sp^2^C is a lone pair-π or n-π^*^ interaction, and that the similarly-favorable interaction of amide sp^3^C with aromatic sp^2^C is a hydrophobic effect and/or a CH-π interaction. The amide sp^2^N-aromatic sp^2^C interaction is marginally unfavorable at 25 °C, as previously determined.^14^

The amide sp^2^C-aromatic sp^2^C interaction (a hydrophobic effect and/or π-π interaction) is favorable, but less so than amide sp^2^O-aromatic sp^2^C and amide sp^3^C-aromatic sp^2^C interactions. The amide sp^2^C-aromatic sp^2^C α-value obtained here (cf. Table 2) is smaller in magnitude than that obtained previously for a combined data set of 85 amide-amide and 20 amide-aromatic μ_23_ value.^15^ This interaction is more challenging to determine than others because of the small amounts of sp^2^C ASA for most amides. Incorporation of the amide-naphthalene data determined here into the previously-analyzed set of amide-amide and amide-aromatic μ_23_ values affects the value of 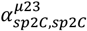. The newly-calculated 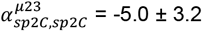 millical mol^-1^ molal^-1^ Å^-4^ for that 109-member set (see SI text and Table S9) agrees within uncertainty with that reported here for the 14-member amide-naphthalene set.

From fitted μ_23_ values at 10 and 45 °C (Table S5), free energy 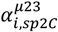 values for interactions of amide compound sp^2^O, sp^2^N, sp^2^C, and sp^3^C atom-types with naphthalene sp^2^C were calculated at these temperatures using Eq. 10. These are summarized together with 25 °C 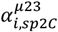 values in the bar graphs of Figure 3A. The favorable interaction of amide sp^2^O with naphthalene sp^2^C becomes less favorable (less negative 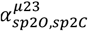) with increasing temperature, demonstrating that the entropic contribution to the favorable sp^2^O-sp^2^C interaction is negative and hence unfavorable. Since the free energy contribution of this amide sp^2^O -naphthalene sp^2^C interaction is favorable 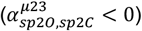, its enthalpic component must be very favorable 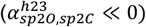.

In contrast to the amide sp^2^O-naphthalene sp^2^C interaction, 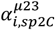 values for interactions of amide sp^2^C and methyl/methylene sp^3^C with naphthalene sp^2^C become more favorable at higher temperature, demonstrating that the entropic contribution to the these interactions is positive and hence favorable. Interactions of amide sp^2^N with naphthalene sp^2^C, slightly unfavorable at 10 °C, also appear to become less unfavorable with increasing temperature, though the uncertainty is large, indicating that here also the intrinsic entropic contribution is positive 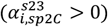. At 45 °C this sp^2^N -sp^2^C interaction appears to be slightly favorable.

### Dissection of Interaction Enthalpies (h_23_) and Entropies (s_23_) Into Atom-atom Contributions

Here we propose and test a dissection of h_23_ and s_23_ values for interactions of 14 amides with naphthalene in order to obtain atom-atom enthalpic and entropic information about interactions of the various unified atoms of the amide compounds investigated (sp^2^O, sp^2^N, sp^2^C, sp^3^C) with naphthalene sp^2^C unified atoms. By analogy with the dissection of 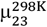 values (Eq. 10), we propose Eqs. 11-12 to interpret 298 s_23_ and h_23_.

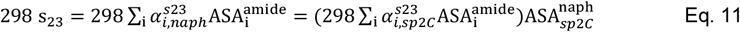

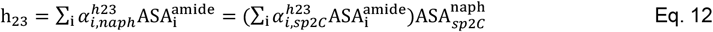

Eqs. 11 and 12 provide both one-way 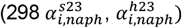 and two-way 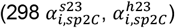 entropic and enthalpic contributions to the corresponding free energy α-values 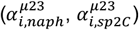 for each amide atom – naphthalene and amide atom – naphthalene atom interaction, where 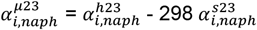 and 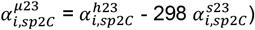. As with Eq. 10, the assumption of additivity is tested in the analysis because the size of the experimental data sets (14 s_23_, 14 h_23_ values) is much greater than the number of fitting parameters (four different 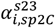, four different 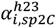). Enthalpic 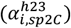 and entropic 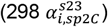 contributions to each 25 °C 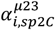 value obtained by application of Eqs. 11-12 are summarized in the bar graphs of Figure 3B and in Table 2.

Figure 3B reveals the very different thermodynamic signature of the amide sp^2^O-aromatic sp^2^C interaction than those of the other atom-atom interactions. The sp^2^O-sp^2^C interaction is strongly enthalpy-driven 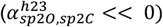, with a large opposing unfavorable entropic contribution 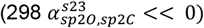. This provides thermodynamic support for a direct chemical interaction of amide sp^2^O atoms with aromatic sp^2^C atoms (e.g. a lone pair-π^20-22^ or n-π^*18,23-25^ interaction).

On the other hand, the interaction of methyl/methylene sp^3^C with naphthalene sp^2^C is entropy-driven 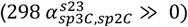, with a large opposing unfavorable enthalpic contribution 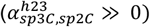. This is the thermodynamic signature of the hydrophobic effect,^26-29^ and indicates that the driving force for this sp^3^C-sp^2^C interaction is the favorable result of removing hydrocarbon surface from exposure to water.

Figure 3B also indicates why the free energy α-value 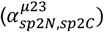 for interaction of amide sp^2^N with aromatic sp^2^C is significantly less favorable than 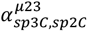 for interaction of methyl/methylene sp^3^C with aromatic sp^2^C. The entropic term is similarly favorable for both these interactions, but is completely compensated by an unfavorable enthalpic term for the amide sp^2^N-aromatic sp^2^C interaction. These sp^2^N-sp^2^C thermodyamics are consistent with a hydrophobic effect (entropic) from reducing the exposure of aromatic sp^2^C surface to water and an offsetting enthalpic cost of disrupting interactions of water with amide sp^2^N.

The opposite signs of the entropic contributions 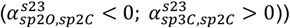) to the intrinsic sp^2^O-sp^2^C and sp^3^C-sp^2^C interaction free energies 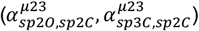 explain the opposite effects of temperature on μ_23_ of interactions of formamide and 1,3-deu with naphthalene in Figure 1. From the values of 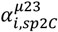 at 25 °C in Table 2 and the ASA information in Table S1 it is straightforward to calculate that the favorable sp^2^O-sp^2^C interaction makes the dominant contribution (∼75%) to the formamide naphthalene μ_23_, while the favorable sp^3^C-sp^2^C interaction makes the dominant contribution (∼90%) to the 1,3-deu-naphthalene μ_23_ (cf. Table S7). The temperature dependences of these dominant atom-atom interactions determine the temperature dependences of the μ_23_ values.

### Enthalpy-Entropy Compensation in Atom-Atom and Solute-Solute Interactions

Strikingly, Figure 3B reveals that all four atom-atom preferential interactions between sp^2^C atoms of naphthalene and sp^3^C, sp^2^O, sp^2^N, and sp^2^C atoms of amide compounds exhibit enthalpy-entropy compensation. Enthalpy-entropy compensation in protein processes has been recently reviewed.^40^ For the atom-atom interactions studied here, enthalpic 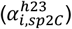 and entropic 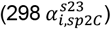 contributions to the intrinsic interaction free energy 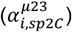 at 25 °C have the same sign and are both much larger in magnitude than 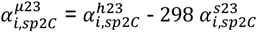. These are relatively weak favorable or neutral atom-atom preferential interactions and do not involve conventional site-binding of the amide compound to naphthalene. The observed enthalpy-entropy compensation may originate in the atom-atom interactions themselves and/or in the interactions with water that are perturbed when an amide atom interacts with a naphthalene atom. For the enthalpically-driven sp^2^O-sp^2^C interaction, magnitudes of 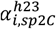 and 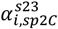 are about ten-fold larger than 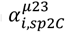 For the equally favorable, entropically-driven sp^3^C-sp^2^C interaction, magnitudes of 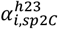 and 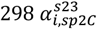 are four and three times larger than 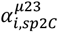. For the less-favorable, entropically-driven sp^2^C-sp^2^C interaction, values of 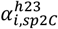 and 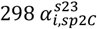 are approximately five and six times larger than 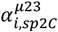, with large uncertainties. For the neutral or slightly unfavorable sp^2^N-sp^2^C interaction, 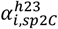 and 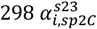 are as large or larger than for sp^3^C-sp^2^C and sp^2^C-sp^2^C interactions and exceed 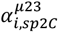 by more than ten-fold.

While all amide atom–naphthalene atom interactions exhibit enthalpy-entropy compensation,Table 1 reveals that only two or three of the fourteen amide molecule – naphthalene molecule interactions investigated here exhibit it to any degree. Interactions of the two diethylureas with naphthalene have positive enthalpic (h_23_) and entropic (298 s_23_) contributions to the interaction free energy 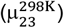 which are at least 1.5 times as large in magnitude as 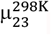. The interaction of formamide with naphthalene has negative h_23_ and 298 s_23_ contributions to 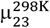 which also are more than 1.5 times as large in magnitude as 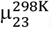 The explanation for these behaviors is provided in Table S7, where the atom-atom 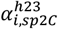 and 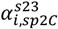 values from Table 2 are used to predict the contributions of each atom-atom interaction to h_23_ and 298 s_23_. For 1,1-deu and 1,3-deu, as a result of the relatively large sp^3^C ASA and relatively small sp^2^O ASA, positive contributions to h_23_ and 298 s_23_ from the sp^3^C-sp^2^C, sp^2^C-sp^2^C, and sp^2^N-sp^2^C interactions sufficiently outweigh the negative contributions to h_23_ and 298 s_23_ from the sp^2^O-sp^2^C interaction to give rise to compensation. Conversely for formamide, which lacks sp^3^C, the negative contributions to h_23_ and 298 s_23_ from the sp^2^O-sp^2^C interaction dominate and give rise to compensation.

Thermodynamic studies of solute effects on protein stability as a function of solute concentration and temperature provide examples of enthalpy-entropy compensation, and of the lack of enthalpy-entropy compensation, at the level of solute molecule – protein molecule preferential interactions. Results for urea, glycine betaine, ethylene glycol and triethylene glycol, obtained from published solute *m*-values,^41-44^ are summarized in Table S10. For unfolding, the solute *m*-value = dΔG°_obs_/dm_solute_ = Δμ = Δh_23_ -TΔs_23_. For solute effects that do not involve site binding, the differences in solute interactions with the unfolded protein vs the folded protein represented by Δμ_23_, Δh_23_ and Δs_23_ are justifiably interpreted as solute interactions with the protein surface exposed in unfolding (i.e. the ΔASA).^2,45^ This ΔASA of unfolding is typically about 2/3 C (mostly sp^3^C) and 1/3 N,O, the majority of which is amide sp^2^N and sp^2^O.

Table S10 reveals that interactions of urea and glycine betaine with the ΔASA of unfolding lacDBD and interactions of urea with the ΔASA of unfolding HPr do not exhibit enthalpy-entropy compensation. Interactions of ethylene glycol and oligomers of polyethylene glycol (e.g.triethylene glycol) with the ΔASA of unfolding SH3 do exhibit enthalpy-entropy compensation (Table S10).^45^ Atom-atom thermodynamic information like that obtained here for interactions of naphthalene sp^2^C atoms with the sp^3^C and sp^2^O, N and C atoms of amide compounds, is needed for interactions of the atoms of these solutes with the sp^3^C, sp^2^N and sp^2^O atoms exposed in unfolding. If the results obtained here for interactions of the atoms of amide compounds with naphthalene atoms (Table 2) are general and the individual atom-atom interactions between the protein ASA exposed in unfolding and urea, GB, EG or PEG polyols in water also exhibit enthalpy-entropy compensation, then these effects must reinforce in the case of EG and small PEGs, but offset one another for urea and GB, like they do for the interaction of urea with naphthalene (Table 1 and Table S7) which does not exhibit enthalpy-entropy compensation.

### Comparison of Experimental Amide-Aromatic 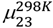, h_23_ and 298 s_23_ with Predictions from Two-Way Atom-Atom α_ij_-Values

Panels A-C of Figure 4 compare observed μ_23_-values at 45, 25 and 10 °C with those predicted from Eq. 10 using the appropriate 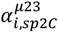 (Tables 2 and S6) and ASA information (Table S1) for each amide. Dotted lines represent perfect agreement of predicted and observed μ_23_-values. Estimated uncertainties in both observed and predicted μ_23_-values are shown; in some cases these are the size of the data points or smaller. Figure 4A-C show that more than 70% of observed μ_23_-values at each temperature agree with predictions from Eq. 10 within the estimated uncertainty.

**Figure 4.**
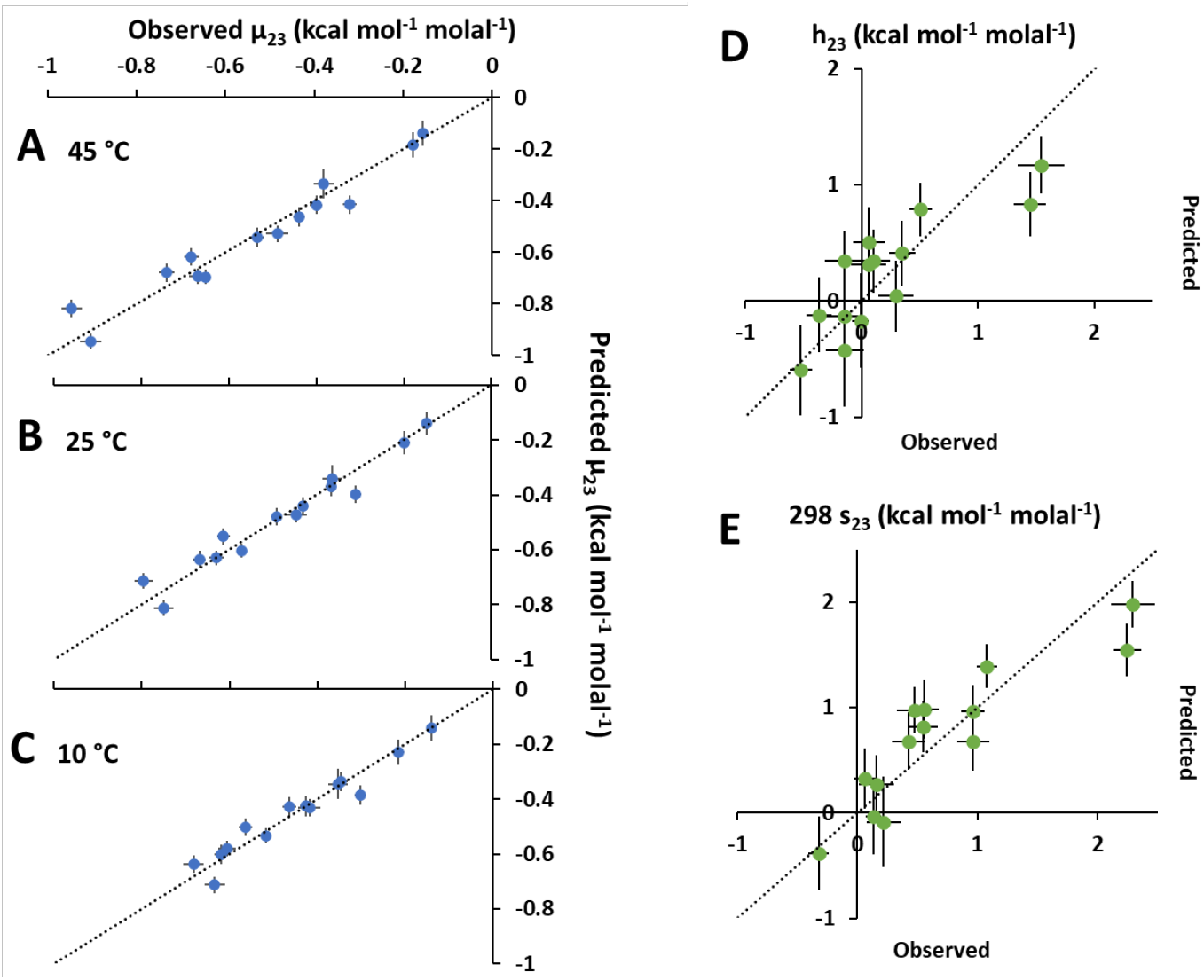
Comparison of predicted and observed μ_23_, h_23_ and 298 s_23_ values. Panels A-C compare predicted and observed μ_23_ values at 45, 25 and 10 °C. Panels D-E compare predicted and observed enthalpic (h_23_) and entropic (298 s_23_) contributions to 25 °C μ_23_ values. Observed μ_23_ values and uncertainties are from Table 1. Predictions of μ_23_ values at each temperature are made using Eq. 10 with the four 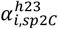 obtained at that temperature (Tables 2, S6) and the solute ASA (Table S1). Predictions of 298 s_23_ and h_23_ are made using Eqs. 11 and 12, the four 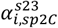 and 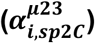 (Table 2) and the solute ASA (Table S1). Uncertainties in all predicted values are propagated as described previously.^11^

Figure 4 D-E compare predicted and observed enthalpic (h_23_) and entropic (298 s_23_) contributions to 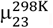. Observed values are reported in Tables S2 and S3, obtained as described above. Predictions are made using Eqs. 11-12, the appropriate 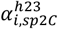 and 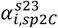 from Table S6, and ASA information (Table S1), and are listed in Tables S2-S3. As expected because of the greater uncertainties in 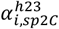 and 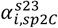 than in 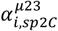, the predicted percentage uncertainties in h_23_ and s_23_ are larger than for 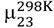. Agreement between predicted and observed values of 298 s_23_ (also h_23_) within the estimated uncertainties is obtained for more than 70% of the amides investigated.

### Intrinsic Strengths of Atom – Atom Interactions of Amides with Amides and Aromatics

The 25 °C amide-naphthalene interaction results (μ_23_ values) reported here supplement and update the set of amide-aromatic μ_23_ values determined previously.^12,14^ Four of the fourteen amide compounds studied here (FAD, mACET, ppa, mad) were not investigated previously, and additional naphthalene solubility data as a function of amide concentration were obtained for the other ten amide compounds. Values of μ_23_ for these fourteen amide-naphthalene interactions were analyzed together with ten μ_23_ values for other amide-aromatic interactions (including amide-anthracene interactions) and eighty-five osmometrically-determined μ_23_ values quantifying interactions of alkylureas and other amides with one another.^14^ This analysis yields an updated set of ten two-way free energy α-values quantifying interactions of sp^2^O, sp^2^N, sp^2^C and sp^3^C atoms of amide and aromatic compounds with one another, shown in Figure 5A and listed in Table S9. In this analysis, as examined and justified previously, amide sp^2^C and aromatic sp^2^C atoms are analyzed together.^12,14^

**Figure 5.**
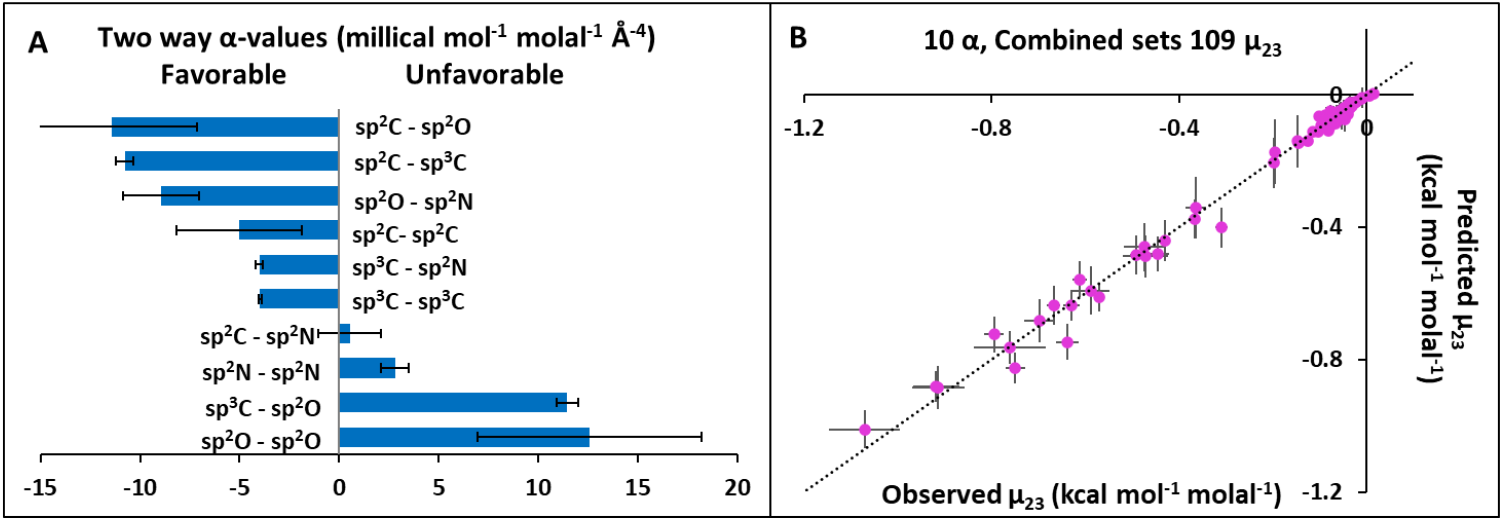
Two-way α_ij_-values quantifying favorable and unfavorable interactions of amide unified atoms with amide and aromatic unified atoms in water. Panel A: Ranking of α_ij_-values obtained from analysis of 85 amide-amide μ_23_ values (VPO, ∼23 °C) and 24 amide-aromatic μ_23_ values (solubility, 25 °C). α_ij_-values and uncertainties are from Table S9. Panel B: Comparison of observed μ_23_ values (listed with uncertainties in Table 1) with those predicted from two-way α_ij_-values (panel A) and ASA information Table S1). See Figure S8D for an expanded scale plot of the amide-amide portion of this figure. Uncertainties in predicted μ_23_ values are propagated as described previously.^11^

All but one of the ten α_ij_-values calculated here (Figure 5) that quantify the different amide atom-amide/aromatic atom interactions are the same within uncertainty as those reported previously.^14^ The exception is 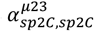, quantifying the intrinsic strength of interactions of amide sp^2^C with amide sp^2^C or aromatic sp^2^C. The value of 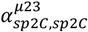 reported in Figure 5 (−5.0 ± 3.1 mcal mol^-1^ molal^-1^ Å^-4^) is the same within uncertainty as that reported in Table 2 from analysis of the amide-naphthalene set of fourteen μ_23_ values (−5.3 ± 1.8 mcal mol^-1^ molal^-1^ Å^-4^), and replaces that determined previously (−11 ± 3 mcal mol^-1^ molal^-1^ Å^-4^).^14^ These values differ by slightly more than the sum of the estimated uncertainties. Because only the three formamides have significant sp^2^C ASA, 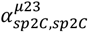 is more difficult to determine than other α_ij_-values. Uncertainties in the other C-C α_ij_-values are much smaller and agreement with those determined previously^14^ is much better. The α_ij_-value for the sp^3^C-sp^2^C interaction is significantly more favorable (−10.8 ± 0.4 mcal mol^-1^ molal^-1^ Å^-4^) than for sp^2^C-sp^2^C, while that for sp^3^C-sp^3^C interaction (−4.0 ± 0.1 mcal mol^-1^ molal^-1^ Å^-4^) is similar in strength to sp^2^C-sp^2^C.

### Investigation of a Chemical Group Alternative (π-π, CH-π) to an Atom-Atom Analysis of Amide-Naphthalene Interactions

A potential alternative to the above atom-atom interpretation of amide-naphthalene interactions is to propose that the amide group and the aromatic ring interact with one another and/or with methyl/methylene sp^3^C atoms primarily as π systems rather than as individual sp^2^C, sp^2^O and sp^2^N atoms. We therefore asked whether the 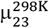, h_23_ and 298 s_23_ results reported here for amide-naphthalene interactions could be analyzed as amide π-aromatic π and methyl/methylene sp^3^C-aromatic π examples of π-π and CH-π interactions, respectively.

Here we test this alternative interpretation by determining the free energy, enthalpy and entropic alpha values for π-π and CH-π interactions and assessing their chemical significance as well as their ability to predict amide-naphthalene μ_23_ values, using Eqs. S5-8. Only two α-values are needed if the amide group is considered as an amide π system with an ASA that is the sum of the amide sp^2^O, sp^2^N and sp^2^C ASA values in Table S1, whereas four α-values are needed when interactions of the sp^2^O, N and C amide atoms are analyzed individually. The two α-value analysis would be warranted if the interaction of the amide group with the aromatic ring primarily involves the π electrons of the amide group and not the individual sp^2^O, N and C atoms of the amide group, which contribute differently to the solvent accessibility of the entire amide group in different amide compounds.

Results of the two-parameter (aromatic π-amide π, CH-aromatic π) analysis are shown in the panels of Figure S7 and in Table S9. Panel A (Figure S7) shows the favorable amide π-aromatic π and CH-aromatic π α-values obtained from analysis of the 14 amide-naphthalene 25 °C μ_23_ values in Table 1 using Eq. S5. Panel A also compares these amide π-aromatic π and CH-aromatic π α-values with the atom-atom α-values (for sp^2^O–sp^2^C, sp^2^N–sp^2^C, sp^2^C–sp^2^C and sp^3^C–sp^2^C interactions; Table 2) obtained from analysis of these μ_23_ values using Eq. 10. The favorable CH-aromatic π α_ij_-value from the two-parameter analysis is almost the same as the sp^3^C–sp^2^C α-value from the four-parameter analysis. Panel A shows that the aromatic π-amide π α-value is in the mid-range of the individual amide atom-aromatic atom α_ij_-values, and much less favorable than the CH-aromatic π α_ij_-value.

Panel B of Figure S7 compares experimentally determined 25 °C amide-naphthalene μ_23_ values with those predicted by the two α_ij_-value (aromatic π-amide π, CH-aromatic π) analysis with those from the atom-atom, four α_ij_-value analysis. Visually it is clear from panel B that, while both analyses predict the observed μ_23_ values quite well, the atom-atom analysis is better than the aromatic π-amide π, CH-π analysis, a conclusion confirmed by the statistical analysis inTable S9. Comparison of root mean square errors (RMSE), calculated by Eq. S10 using differences between observed and predicted 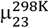 (Table S9), reveals that RMSE is ∼20%larger for the analysis using two α-values. Expanding the 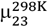 data set to 24 amide-aromatic interactions using anthracene and additional naphthalene results obtained only at 298 K yields the same conclusion. These 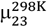 are quite well-fit using two α_ij_-values which are the same within uncertainly as for the 14 amide set, but from both a chemical and statistical perspective the analysis using four α-values is superior (Table S9; Figure S6 vs S8A). Likewise the analysis of μ_23_ values for amide-naphthalene interactions at 10 and 45 °C (Figure S7 panels C, D) reveals that atom-atom α_ij_-values provide a better interpretation both statistically and chemically.

We conclude that the aromatic π-amide π α-values 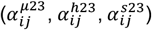 are better interpreted as weighted-average approximations to the atom-atom α-values for the different amides in the data set, rather than as π-π interactions. The ability of the aromatic π-amide π, CH-π analysis to fit the data with this average α_ij_-value for the range of amide groups on the 14 amide compounds investigated shows why the uncertainties in the individual atom-atom α_ij_-values for the amide group are relatively large (Tables 2, S9), because of residual correlations between the atom-atom α_ij_-values for amide sp^2^O, N and C atoms.

Two and four α-value analyses of 298 s_23_ and h_23_ values (Eq. S7-8; Figure S7 panels F,G) give similar results. While predictions of 298 s_23_ and h_23_ using only two α-values (π-π, CH-π, Table S8) are almost as good as those obtained using four atom-atom α-values (Table S6), in all cases the quality of the fit as assessed by RMSE is better using the four atom-atom α-values.

In summary, although the two α-value analysis of these data as π interactions described in this section is moderately successful from a statistical perspective, the four parameter analysis as atom-atom interactions is in all cases (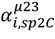at 10, 25 and 45 °C; 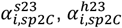) a better statistical fit. Also, the thermodynamics that we determine from both analyses for the very favorable interaction of methyl and methylene portions of the amide compound with naphthalene (entropy driven, enthalpically unfavorable) are more consistent with this being a hydrophobic effect, driven by the entropic effect of releasing local water from these hydrocarbon surfaces when they interact, than what would be expected for a direct CH-π interaction.

We also tested whether the combined 109 member set of amide-amide and amide-aromatic μ_23_ values could be successfully interpreted if interactions with the amide group are analyzed as interactions with the amide π system rather than as interactions with amide sp^2^O, sp^2^N and sp^2^C atoms, using Eqs. S10A and B. Table S9 summarizes the α-values obtained from this analysis and compares RMSE values for 4 and 10-parameter analyses. Figures S8 compares predicted and observed 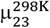 values using these different fitting analyses. Figure S8B shows the full range of the 109 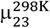 data set, which is dominated by the amide-aromatic portion, while Figure S8C focus on the amide-amide portion of that large 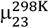 data set. The four parameter analysis predicts a subset of the 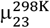 values quite well, as shown in Figure S8B, but statistical analysis (RMSE; Table S9) reveals that it is not as good as the fit involving ten α-values. More importantly, α-values for the combined amide group lack the molecular information provided by individual amide atom α-values, which make good chemical sense as discussed above and previously.^12,14^

In summary, while the reduced parameter fits do quite well, the atom-atom fits are better. Where α-values in the reduced parameter fits have an atom-atom interpretation, like CH-aromatic π as sp^3^C-sp^2^C, these α-values are very similar to the atom-atom α-values. The amide π α-values behave like rough averages of the atom-atom alpha values for amide sp^2^O, sp^2^N and sp^2^C and provides no support for the idea that amide interations are primarily π system interactions as opposed to individual atom-atom interactions. Atom-atom α-values provide a better analysis of all these amide interaction data, both statistically and chemically.

## Conclusions

We conclude from measurements of effects of 14 amide compounds on solubility of naphthalene at 10, 25 and 45 °C that the thermodynamics of amide compound-naphthalene interactions (μ_23_, h_23_, s_23_) are well-analyzed by the ASA-based approach previously applied to μ_23_ values for amide interactions at 25 °C.^14^ This analysis makes and tests the assumption of additivity of the different atom-atom interactions to molecule-molecule interactions (μ_23_, h_23_, s_23_). In this analysis, the intrinsic contribution of each atom-atom interaction (expressed per unit of accessible surface area of each atom) is weighted by the product of ASA of these interacting atoms on the two molecules. This use of surface area as the fundamental variable is justified in previous SPM analyses of solute-protein interactions^46^ and solute and salt effects on surface tension^47-49^. For amide-amide and amide-aromatic interactions, use of ASA as the fundamental variable was shown to be superior to an analysis based only on the number of each type of atom and not its accessibility.^12,14^

From the temperature dependences of μ_23_ values for these 14 amide compounds with naphthalene, we dissect μ_23_ values into their enthalpic (h_23_) and entropic (298 s_23_) contributions at 25 °C. For the series of alkylureas and formamides, μ_23_, h_23_ and 298 s_23_ all vary systematically with amide composition. Analysis of these results yields enthalpic 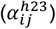 and entropic 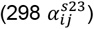 contributions to intrinsic strengths of interactions 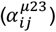 of amide sp^2^O, sp^2^N, sp^2^C and sp^3^C unified atoms with aromatic sp^2^C atoms. These enthalpic and entropic contributions have the same sign and are much larger in magnitude than the interaction free energy itself. The amide sp^2^O-aromatic sp^2^C interaction is enthalpy-driven and entropically unfavorable, consistent with direct chemical interaction (e.g. lone pair-π). On the other hand, amide sp^3^C- and sp^2^C-aromatic sp^2^C interactions are entropy-driven and enthalpically unfavorable, consistent with hydrophobic effects, and the amide sp^2^N-aromatic sp^2^C interaction is entropically favorable, as if hydrophobic, with an unfavorable enthalpic term of equal magnitude so the net free energy of this interaction is zero within uncertainty. These findings will be relevant to interpret interactions involving π-bonded sp^2^ atoms in protein processes.

## Methods

### Experimental Methods

Solubility assays are performed and analyzed as previously described.^10-12^ Additional details of these assays, and information about the naphthalene and amide samples used in this research, are given in SI Methods.

## Acknowledgements

We thank Alana Bordeaux and Gavin Frings for their experimental contributions, and acknowledge NIH grant R35 GM 118100 and the University of Wisconsin-Madison for support of this research.

