## Supplemental information for "Quantifying Amide-Aromatic Interactions at Molecular and Atomic Levels: Experimentally-determined Enthalpic and Entropic Contributions to Interactions of Amide sp^2^O, N, C and sp^3^C Unified Atoms with Naphthalene sp^2^C Atoms in Water"

**Chemicals**

Acetamide (purity ≥ 99.0%), 1,1-diethylurea (97%), 1,1-dimethylurea (99%), 1,3-diethylurea (97%), 1,3-dimethylurea (99%), dimethylformamide (99.8%), ethylurea (97%), methylacetamide (≥99%), methylformamide (99%), and naphthalene (≥ 99%) were obtained from Sigma-Aldrich. Malonamide (98%) was obtained from Alfa Aesar. Propionamide was obtained from Alfa Aesar (98%) and TCI America (98.0%). Methylurea (97%) was obtained from Acros Organics. Formamide (≥ 99.5%) was obtained from Fisher. Urea (99%) was obtained from VWR. All solutions were prepared with 18 MΩ water purified with a Barnstead E-pure system (Thermo Fisher Scientific).

**Preparation of Samples**

Naphthalene as received from the supplier dissolves very slowly. To increase the dissolving rate, solid samples were ground with a mortar and pestle and then placed in a beaker, covered with water, and homogenized for five minutes at 20,000 rpm using a OMNI TH Tissue Homogenizer with a 7 mm stainless steel probe, similar to a previous approach.[1] Naphthalene samples were then dried at room temperature in a fume hood for several days before storing in a desiccator with Drierite (anhydrous calcium sulfate) to absorb any remaining water. In the desiccator, water is found to evaporate from the solid naphthalene faster than the rate of sublimation of naphthalene, resulting in a 1.0% reduction in mass of the naphthalene sample in the first 24 hours (from evaporation of water and naphthalene) followed by a 0.65 ± 0.05% per day reduction in mass for the next week, which we interpret as from evaporation of naphthalene only. Experimental results obtained with naphthalene samples dried for various times in excess of one day exhibited no systematic differences.

All solutions are prepared as previously described.[2-4] Briefly, for each trial a set of 12 samples is prepared with amide concentrations typically in the range 0 - 2 m, in 15 mL polypropylene centrifuge tubes. An excess of naphthalene is added, and the samples are tightly capped, vortexed, and placed in a temperature-regulated (± 0.1 °C) water bath or air shaker (Grant Scientific OLS AquaPro or OLS200, Thermo Scientific MaxQ 4000). Naphthalene adsorbs to polypropylene, but this causes no problems because of the excess of solid naphthalene.

At 45 °C, solubility equilibrium is attained in 3 to 5 days. Stable determinations of the amount of naphthalene dissolved at 45 °C were obtained for at least 10 days after this, and most determinations were made after one week of shaking. At 25 °C, solubility equilibrium is attained in 5 to 7 days. Stable determinations of the amount of naphthalene dissolved at 25 °C were obtained for at least two weeks, and most determinations were made after one week of shaking. At 10 °C, two weeks is required to reach solubility equilibrium. To reduce the time required, samples were incubated at 45 °C for two days, and then transferred to 10 °C. Solubility equilibrium at 10 °C was attained after an additional 3-7 days of shaking and determinations of the amount dissolved were stable over at least a two week period after that.

In order to determine naphthalene concentration by absorbance, samples are diluted by mass so that peak absorbance at 276 nm is less than 1. Dilutions are prepared in glass centrifuge tubes because naphthalene adsorbs to polypropylene tubes, reducing its concentration. Any suspended particles of homogenized naphthalene in these solutions are eliminated by centrifugation at 8,000 rpm for 10 minutes before determining the absorbance. Absorbances of diluted samples in 3 mL quartz cuvettes are determined relative to a matched blank cuvette at 276 nm in an Agilent Cary 300 Spectrophotometer thermostatted at 25 °C. None of the amide compounds investigated here absorbs appreciably at this wavelength.

### Water-Accessible Surface Area

The water accessible surface area (ASA) of acetamide was calculated from the structure generated by CACTUS SMILES (http://cactus.nci.nih.gov/ translate/) using Surface Racer,[5] the Richards set of van der Waals radii[6] and a 1.4 Å radius for water as previously described.[2,3] CACTUS was used previously to generate structures of other amides without structural data. Where comparisons were possible, ASA results from CACTUS structures agreed within 10% (comparable to the experimental uncertainty in $\mu_{23}$ values) with those obtained from BMRB files.[2,3] ASA of all other amide compounds investigated here as well as naphthalene and anthracene were determined and dissected into contributions from the different hybridization states of unified O, N and C atoms previously.[3] These include amide $sp^2O$, $sp^2N$ and $sp^2C$ and of methyl/methylene $sp^3C$ of amide compounds, and aromatic $sp^2C$ of naphthalene and anthracene, as listed in Table S1. Intrinsic strengths of interaction (i.e. per unit ASA of each type of atom) of amide $sp^2C$ and aromatic $sp^2C$ with other atom types were found to be similar (about 30% more favorable for aromatic $sp^2C$) and so are analyzed together, as discussed previously.[3,7]

### Determination of the thermodynamics of amide compound-naphthalene preferential interactions ($\mu_{23}$, $h_{23}$, and $s_{23}$) from solubility data

Initial steps of analysis of solubility data to obtain chemical potential derivatives $(\partial\mu_2/\partial m_3)_{P,T,m2} = \mu_{23}$ quantifying preferential interactions of amides (component 3) with naphthalene (component 2) at each temperature are performed as previously described.[2-4] Absorbances are corrected for dilution and converted from a molar to a molal concentration scale as previously described.[2] The logarithm of the corrected absorbance is plotted vs amide concentration for data in one experiment and fitted to obtain a best-fit intercept, which is used to normalize data from that experiment and obtain values of $\left(m_2^{ss,m3}/m_2^{ss,0}\right)$, the ratio of the naphthalene concentration in the saturated solution at amide concentration $m_3$ ($m_2^{ss,m3}$) relative to that in the absence of amide ($m_2^{ss,0}$). For each amide compound investigated, values of $\ln(m_2^{ss,m3}/m_2^{ss,0})$ are plotted as a function of amide concentration $m_3$ and fit to a quadratic function of $m_3$ with an intercept of zero. Standard deviations (SD) are obtained for the large sets of two component naphthalene solubility data (determinations from all amide series for $m_3 = 0$ at each temperature) and are used to determine outliers in three-component data. Approximately 10% of three-component data points deviate from the best fit curve by more than ± 2 SD and were removed before analysis of naphthalene $\Delta G^o_{obs,tr}$ as a function of amide concentration and temperature, described below.

**Details of analysis of naphthalene $\Delta G^o_{obs,tr}$ to obtain $\mu_{23}$, $s_{23}$ and $h_{23}$ of each amide-naphthalene interaction**

Because the thermodynamic activity of naphthalene in a saturated solution at a specified temperature is independent of amide concentration, the observed standard free energy of transfer ($\Delta G^o_{tr}$; Eq. 1) of naphthalene (component 2) from water to an amide (component 3) solution at molal concentration $m_3$ is interpreted in Eq. S1 in terms of the ratio $\gamma_{2,m3}/\gamma_{2,0}$ where $\gamma_{2,m3}$ and $\gamma_{2,0}$ are molal scale activity coefficients of naphthalene at amide concentrations $m_3$ and at $m_3 = 0$.

$$\Delta G^o_{obs,tr} = RT \ln\left(\frac{\gamma_{2,m3}}{\gamma_{2,0}}\right) \qquad \text{Eq. S1}$$

Because the solubility of naphthalene is very low and the amide concentration is in excess, $\gamma_{2,m3}$ is only a function of $m_3$ and T and not a function of $m_2$.

The dependence of $\gamma_{2,m3}$ on $m_3$ determines both $\mu_{23}$ and $\mu_{233}$.

$$\mu_{23} = (\partial\mu_2/\partial m_3)_{T,P,m_2} = RT\left(\partial ln\gamma_{2,m3}/\partial m_3\right)_{T,P,m_2} = \left(\partial\Delta G^o_{obs,tr}/\partial m_3\right)_{T,P,m_2} \qquad \text{Eq. S2}$$

$$\mu_{233} = (\partial^2\mu_2/\partial m_3^2)_{T,P,m_2} = RT\,(\partial^2 ln\gamma_{2,m3}/\partial m_3^2)_{T,P,m_2} = (\partial^2\Delta G^o_{obs,tr}/\partial m_3^2)_{T,P,m_2} \quad \text{Eq. S3}$$

In the text, values of $\Delta G^o_{obs,tr}$ for each amide as a function of $m_3$ and T, calculated from the solubility data using Eq. 4, are fit globally to Eq. 8 using Igor Pro 5.05A to obtain $\mu_{23}^{298K}$, $s_{23}$ and $\mu_{233}$ where $s_{23}$ is assumed to be independent of temperature and $m_3$. Then $h_{23}$ is obtained from $\mu_{23}^{298K}$ and $s_{23}$ using Eq. 9. An alternative approach is to analyze values of $\Delta G^o_{obs,tr}$/T for each amide as a function of $m_3$ and 1/T using the relationship:

$$\Delta G^o_{obs,tr}/T = (\mu_{23}^{298K}/298)m_3 - (h_{23}/298)m_3(1 - 298/T) + 0.5\,(\mu_{233}/298)m_3^2 \quad \text{Eq. S4}$$

Fitting to Eq. S4 yields $h_{23}$, assumed to be independent of temperature and $m_3$. From $h_{23}$ and $\mu_{23}^{298K}$, $s_{23}$ is obtained using Eq. 9. Values of $h_{23}$ (and of 298 $s_{23}$), listed with their propagated uncertainty[2] in Tables S2 and S3, do not differ significantly for the two analyses. Global fits including a non-zero heat capacity of the amide-naphthalene interaction, so that $h_{23}$ and $s_{23}$ vary with temperature, were also tested but did not improve the fit.

**Comparison of values of $\mu_{23}$ and $\alpha_{i,sp2C}^{\mu 23}$ at 298 K with published results**

Values of $\mu_{23}$ for interactions of fourteen amide compounds with naphthalene at three temperatures (10, 25 and 45 °C) were determined in the research reported here. Values of $\mu_{23}^{298K}$ for interactions of ten of these amide compounds with naphthalene at 25 °C were reported previously.[3] Solubility data as a function of $m_3$ for nine of the ten previously-investigated amides agree within uncertainty with the published results and were analyzed together to obtain the $\mu_{23}^{298K}$ values reported here. For seven of these nine amides, these $\mu_{23}^{298K}$ values are the same within uncertainty as those previously reported.[3] For two other amides (ethylurea, acetamide), $\mu_{23}^{298K}$ values reported here differ from those previously reported by somewhat more than the combined experimental uncertainty. For methylformamide, four independent solubility series agree well with each other but differ systematically from the published result[3] and we report $\mu_{23}^{298K}$ obtained from these four determinations here.

These changes in the set of amide-aromatic $\mu_{23}^{298K}$ values affect $\alpha_{sp2C,sp2C}^{\mu23}$, the intrinsic strength of the amide $sp^2C$–aromatic $sp^2C$ interaction, reducing the magnitude of this favorable interaction by approximately 50% from -11 ± 3 (from analysis of 85 amide-amide $\mu_{23}$ and the original set of 20 amide-aromatic $\mu_{23}$)[7] to -5 ± 3 from analysis of 85 amide-amide $\mu_{23}$ and the current set of 24 amide-aromatic $\mu_{23}$ (cf. Table S9). This change in $\alpha_{sp2C,sp2C}^{\mu23}$ is marginally significant given the combined uncertainties. The same $\alpha_{sp2C,sp2C}^{\mu23}$ (-5 ± 2) is obtained from analysis of the set of 14 amide-naphthalene $\mu_{23}^{298K}$ values reported here. Analysis of the larger set of 24 amide-aromatic $\mu_{23}^{298K}$ values yields a similar $\alpha_{sp2C,sp2C}^{\mu23}$ with a larger uncertainty. The other nine two-way α-values determined for amide-aromatic and amide-amide interactions (Table S9) are not significantly affected by these changes in the $\mu_{23}^{298K}$ data set.

**Tests of robustness of results of the ASA-based thermodynamic analysis; Effects from removal of one solute on $\alpha_{ij}$-values and on entropic and enthalpic contribution**

As a test of whether any one amide-naphthalene interaction has an unusually large impact on determination of the $\alpha_{ij}$-values, $\mu_{23}$ values at the three temperatures for one of the 14 amide compounds were deleted and $\mu_{23}$ values for the remaining 13 amides at the three temperatures were analyzed to obtain the four $\alpha_{ij}$-values. Similar analyses were performed with $s_{23}$ and $h_{23}$ values. Results are shown in the bar graphs of Figures S3-S5.

Figure S3, reporting the analysis of subsets of $\mu_{23}$ values, shows that the 25 °C value of $\alpha_{sp3C,sp2C}^{\mu23}$, the intrinsic strength of the $sp^3C$-$sp^2C$ interaction, is completely insensitive to removal of $\mu_{23}$ values for any one of the 14 amide compounds. This free energy $\alpha_{ij}$-value is very robust because $sp^3C$ ASA on the 14 amide compounds varies widely and is uncorrelated with amide

ASA, as shown in Table S1. Figure S3 also shows that the three $\alpha_{i,sp2C}^{\mu23}$ values for interactions of amide $sp^2O$, $sp^2N$ and $sp^2C$ with aromatic $sp^2C$ are not quite as robust as $\alpha_{sp3C,sp2C}^{\mu23}$. This is probably because of residual correlations between amounts of amide $sp^2O$, $sp^2N$ and $sp^2C$ ASA in subsets of these amide compounds, since these atom-types are present in a 1:1:1 ratio for all amides investigated except urea. Gratifyingly, these three amide atom-aromatic atom free energy $\alpha_{ij}$-values are insensitive (i.e. don't differ by more than the combined experimental uncertainties) to deletion of any of 12 amide compounds. However, deletion of acetamide results in an $\alpha_{sp2O,sp2C}^{\mu23}$ that is significantly more favorable and $\alpha_{sp2N,sp2C}^{\mu23}$ and $\alpha_{sp2C,sp2C}^{\mu23}$ that are significantly more unfavorable than those for the entire 14 amide dataset. The opposite situation is observed if 1,3-diethylurea is deleted. In this case $\alpha_{sp2O,sp2C}^{\mu23}$ becomes significantly less favorable and $\alpha_{sp2N,sp2C}^{\mu23}$ and $\alpha_{sp2C,sp2C}^{\mu23}$ become significantly more favorable than those for the entire 14 amide dataset. Inclusion of $\mu_{23}$ values for both acetamide and 1,3-diethylurea makes all three of these amide-aromatic free energy $\alpha_{ij}$-values much more robust, so they are insensitive to removal of $\mu_{23}$ values for any one of the other 12 amide compounds in the dataset.

Figures S4 and S5, reporting the analysis of subsets of values of 298 $s_{23}$ and $h_{23}$, reveal that the entropic ($298\ \alpha_{\mathrm{sp3C,sp2C}}^{\mathrm{s23}}$) and enthalpic ($\alpha_{\mathrm{sp3C,sp2C}}^{\mathrm{h23}}$) contributions to the intrinsic strength of the $sp^3C$-$sp^2C$ interaction, are (like $\alpha_{\mathrm{sp3C,sp2C}}^{\mu23}$ in Figure S3) completely insensitive to removal of the $s_{23}$ or $h_{23}$ value for any one of the 14 amide compounds. Figures S4 and S5 also show that the $298\ \alpha_{\mathrm{i,sp2C}}^{\mathrm{s23}}$ and $\alpha_{\mathrm{i,sp2C}}^{\mathrm{h23}}$ values for interactions of amide $sp^2O$, $sp^2N$ and $sp^2C$ with aromatic $sp^2C$ are as robust as $298\ \alpha_{\mathrm{sp3C,sp2C}}^{\mathrm{s23}}$ and $\alpha_{\mathrm{sp3C,sp2C}}^{\mathrm{h23}}$, respectively. Values of $298\ \alpha_{\mathrm{sp3C,sp2C}}^{\mathrm{s23}}$ and $\alpha_{\mathrm{sp3C,sp2C}}^{\mathrm{h23}}$ are insensitive (i.e. don't differ by more than the combined uncertainties) to the deletion of any of 13 amide compounds. However, deletion of methylacetamide affects entropic and enthalpic contributions to $\alpha_{ij}$-values for amide atom -

naphthalene atom interactions significantly. Deletion of methylacetamide results in a $298\ \alpha^{s23}_{sp2O,sp2C}$ that is significantly less unfavorable and $298\ \alpha^{s23}_{sp2N,sp2C}$ and $298\ \alpha^{s23}_{sp2C,sp2C}$ that are significantly less favorable than those for the entire 14-amide dataset. Figure S4 shows that $298\ \alpha^{s23}_{sp2C,sp2C}$ becomes negative if methylacetamide is deleted, while it is positive as expected for a hydrophobic amide $sp^2$C-aromatic $sp^2$C interaction for the 14 amide set and all other single-amide deletions. As expected from the observation that deletion of methylacetamide has no significant effect on the free energy $\alpha_{ij}$-values (Figure S3), deletion of methylacetamide affects the enthalpic contributions to these $\alpha_{ij}$-values in the opposite direction to its effects on the entropic $\alpha_{ij}$-values, as shown in Figure S5.

**Equations for analysis of amide-naphthalene interaction data ($\mu_{23}, h_{23}, s_{23}$; Table 1) as π-π and CH–π interactions**

If interactions of amide compounds with naphthalene can be described as π-π and CH–π interactions, with free energy $\alpha_{ij}$-values $\alpha^{\mu23}_{\pi-\pi}$ and $\alpha^{\mu23}_{CH-\pi}$, Eq. 10 for $\mu_{23}$ becomes

$$\mu_{23} = (\alpha^{\mu23}_{\pi-\pi}\,\mathrm{ASA}^{\mathrm{amide}}_{\pi} + \alpha^{\mu23}_{\mathrm{CH}-\pi}\,\mathrm{ASA}^{\mathrm{amide}}_{\mathrm{CH}})\mathrm{ASA}^{\mathrm{naph}}_{\pi} \qquad \text{Eq. S5}$$

where

$$\mathrm{ASA}^{\mathrm{amide}}_{\pi} = \mathrm{ASA}^{\mathrm{amide}}_{\mathrm{sp2O}} + \mathrm{ASA}^{\mathrm{amide}}_{\mathrm{sp2N}} + \mathrm{ASA}^{\mathrm{amide}}_{\mathrm{sp2C}} \quad \text{and} \quad \mathrm{ASA}^{\mathrm{amide}}_{\mathrm{CH}} = \mathrm{ASA}^{\mathrm{amide}}_{\mathrm{sp3C}} \qquad \text{Eq. S6}$$

Fitted values of $\alpha^{\mu23}_{\pi-\pi}$ and $\alpha^{\mu23}_{CH-\pi}$ were obtained from analysis of the set of 14 amide-naphthalene 25 °C $\mu_{23}$ values (results listed in Table S8). A larger set including these 14 amide-naphthalene $\mu_{23}$ values and 10 previously-published $\mu_{23}$ values for other amide-aromatic interactions[3] is also analyzed and results are listed in Table S9. Comparison of the four $\alpha_{ij}$-value fits to the two $\alpha_{ij}$-value fits are shown in Figure S7. Analogous fits to the 14 amide-naphthalene 298 $s_{23}$ values and $h_{23}$ values (Tables S2, S3) were performed to obtain entropic and enthalpic contributions to these $\alpha^{\mu23}_{\pi-\pi}$ and $\alpha^{\mu23}_{CH-\pi}$ values.

$$298\ s_{23} = (298\ \alpha^{s23}_{\pi-\pi}\,\mathrm{ASA}^{\mathrm{amide}}_{\pi} + 298\ \alpha^{s23}_{\mathrm{CH}-\pi}\,\mathrm{ASA}^{\mathrm{amide}}_{\mathrm{CH}})\mathrm{ASA}^{\mathrm{naph}}_{\pi} \qquad \text{Eq. S7}$$

$$h_{23} = (\alpha^{h23}_{\pi-\pi}\ ASA^{amide}_{\pi} + \alpha^{h23}_{CH-\pi}\ ASA^{amide}_{CH})ASA^{naph}_{\pi} \qquad \text{Eq. S8}$$

These π–π and CH-π contributions to $\alpha_{ij}$-values are listed in Table S8 and shown in the bar graph in Figure S7E.

**Equations for analyses of amide-amide and amide-aromatic interaction data ($\mu_{23}$) in terms of π-π and CH–π interactions**

A similar analysis based on interactions of amide and aromatic π systems instead of individual atom-atom interactions was performed with the combined set of 109 amide-aromatic and amide-amide $\mu_{23}$ values from ref. 7 and this work, using four $\alpha_{ij}$-values ($\alpha^{\mu23}_{sp3C,sp3C}$, $\alpha^{\mu23}_{sp3C,am\,\pi}$, $\alpha^{\mu23}_{sp3C,aro\,\pi}$, $\alpha^{\mu23}_{\pi,\pi}$). This fit introduces individual $\alpha_{ij}$-values ($\alpha^{\mu23}_{sp3C,am\,\pi}$, $\alpha^{\mu23}_{sp3C,aro\,\pi}$) for interactions of amide $sp^3C$ with amide and aromatic π systems (i.e. different CH-π $\alpha_{ij}$-values for interactions of $sp^3C$ with amide and aromatic π). A common $\alpha_{ij}$-value ($\alpha^{\mu23}_{\pi,\pi}$) is assumed applicable to amide π-amide π and amide π-aromatic π interactions.

$$\mu_{23} = \alpha^{\mu23}_{sp3C,sp3C} ASA^{amide2}_{sp3C} ASA^{amide3}_{sp3C} + \alpha^{\mu23}_{sp3C,am\,\pi}\left(ASA^{amide2}_{sp3C} ASA^{amide3}_{\pi} + ASA^{amide2}_{\pi} ASA^{amide3}_{sp3C}\right)$$

$$+\ \alpha^{\mu23}_{\pi,\pi}(ASA^{amide2}_{\pi} ASA^{amide3}_{\pi} + ASA^{amide}_{\pi} ASA^{aro}_{\pi}) +\ \alpha^{\mu23}_{sp3C,aro\,\pi} ASA^{amide}_{sp3C} ASA^{aro}_{\pi} \qquad \text{Eq. S9}$$

where $ASA^{aro}_{\pi}$ is the ASA of naphthalene or anthracene and $ASA^{amide}_{\pi}$ is defined in Eq. S6.

An alternative four $\alpha_{ij}$-value fit to the 109 member set (not shown), using a common $\alpha_{ij}$-value for interactions of amide $sp^3C$ with both amide and aromatic π systems ($\alpha^{\mu23}_{sp3C,am/aro\,\pi}$) and individual $\alpha_{ij}$-values for amide-π–amide π ($\alpha^{\mu23}_{am\,\pi,am\,\pi}$) and amide π–aro π ($\alpha^{\mu23}_{am\,\pi,aro\,\pi}$) failed on both statistical and chemical grounds, yielding an unfavorable $sp^3C$-$sp^3C$ $\alpha_{ij}$-value ($\alpha^{\mu23}_{sp3C,sp3C} > 0$) and a larger root mean square error (RMSE), calculated as described below.

### Calculation of root mean square errors (RMSE)

To assess the quality of all fits to all data sets analyzed here we calculate the root mean square error (RMSE),[8] defined as the square root of the quotient of the SSE (the sum of squares of differences between observed and predicted $\mu_{23}$ values) divided by the DOF (the degrees of freedom, defined as the difference n – k where n is the number of values of $\mu_{23}$ , $h_{23}$ or 298 $s_{23}$ being analyzed and k is the number of $\alpha_{ij}$-values determined from the fit.

For $\mu_{23}$

$$\mathrm{RMSE} = \sqrt{\frac{\mathrm{SSE}}{DOF}} = \sqrt{\frac{\sum_1^n(\mu_{23,i}^{obs} - \mu_{23,i}^{pred})^2}{n-k}} \qquad \text{Eq. S10}$$

In Eq. S10, $\mu_{23,i}^{obs}$ and $\mu_{23,i}^{pred}$ are observed and predicted values of $\mu_{23}$ for member i of the set being analyzed. In these analyses, n ranged from 14 to 109 $\mu_{23}$ values and k ranged from 2 to 10 $\alpha_{ij}$-values. Definitions of RMSE for $h_{23}$ and 298 $s_{23}$ are analogous to that in Eq. S10 for $\mu_{23}$. For different fits (i.e. different choices of k) to a given data set (i.e. a specified value of n), the smaller the RMSE the better the fit. Values of RMSE for different fits and/or data sets are tabulated in Tables 2, S6, S8 and S9.

**Table S1. Water-accessible surface areas (ASA) of amide and aromatic compounds [a]**

| Solute | ASA, Å² | | | |
|---|---|---|---|---|
| | Aliphatic $sp^3C$ | Amide/Aromatic $sp^2C$ | Amide $sp^2O$ | Amide $sp^2N$ |
| urea | 0 | 7.2 | 47.9 | 130 |
| methylurea | 88.4 | 6.5 | 38.3 | 87.5 |
| ethylurea | 125 | 6.5 | 38.3 | 82.0 |
| 1,1-dimethylurea | 149 | 6.2 | 38.3 | 54.5 |
| 1,3-dimethylurea | 177 | 5.8 | 28.7 | 44.9 |
| 1,1-diethylurea | 209 | 3.7 | 35.7 | 50.0 |
| 1,3-diethylurea | 249 | 5.8 | 28.7 | 33.8 |
| malonamide | 48.5 | 8.5 | 65.7 | 123 |
| propionamide | 125 | 4.3 | 36.8 | 61.6 |
| formamide | 0 | 40.2 | 51.3 | 70.8 |
| methylformamide | 88.4 | 39.5 | 41.7 | 27.7 |
| dimethylformamide | 158 | 30.7 | 41.7 | 0.763 |
| acetamide | 89.7 | 4.3 | 44.9 | 61.6 |
| methylacetamide | 178 | 3.6 | 35.3 | 19.0 |
| naphthalene | 0 | 273 | 0 | 0 |
| anthracene | 0 | 334 | 0 | 0 |

[a] Previously published[3] except acetamide, calculated as described in SI Methods.

**Table S2. Comparison of predicted 298 $s_{23}$ (kcal mol$^{-1}$ molal$^{-1}$) from $\alpha_{i,sp2C}^{s23}$ with results of global fitting of $\Delta G_{tr}^{o}$ data**

| Solute | Fitted (Eq. 8) [a,b] | Fitted (Eq. S4) [c] | Predicted (Eq. 11) [b,d] |
|---|---|---|---|
| **urea** | 0.14 ± 0.06 | 0.03 ± 0.14 | -0.03 ± 0.36 |
| **methylurea** | 0.43 ± 0.14 | 0.53 ± 0.12 | 0.68 ± 0.28 |
| **ethylurea** | 0.56 ± 0.13 | 0.61 ± 0.13 | 0.98 ± 0.27 |
| **1,1-dmu** | 0.96 ± 0.10 | 1.07 ± 0.10 | 0.96 ± 0.25 |
| **1,3-dmu** | 1.08 ± 0.09 | 1.16 ± 0.09 | 1.39 ± 0.21 |
| **1,1-deu** | 2.24 ± 0.12 | 2.34 ± 0.12 | 1.54 ± 0.25 |
| **1,3-deu** | 2.29 ± 0.18 | 2.46 ± 0.18 | 1.98 ± 0.22 |
| **malonamide** | 0.22 ± 0.14 | 0.28 ± 0.14 | -0.09 ± 0.43 |
| **propionamide** | 0.55 ± 0.12 | 0.65 ± 0.12 | 0.82 ± 0.24 |
| **formamide** | -0.32 ± 0.09 | -0.29 ± 0.09 | -0.38 ± 0.35 |
| **methylFAD** | 0.07 ± 0.09 | 0.11 ± 0.10 | 0.32 ± 0.29 |
| **dimethylFAD** | 0.96 ± 0.13 | 1.07 ± 0.14 | 0.68 ± 0.27 |
| **acetamide** | 0.16 ± 0.14 | 0.16 ± 0.14 | 0.27 ± 0.28 |
| **methylacetamide** | 0.48 ± 0.14 | 0.61 ± 0.16 | 0.98 ± 0.22 |

[a] Obtained by global fitting to Eq. 8 by Igor Pro 5.0.5A. Uncertainties are twice the fitting error.
[b] Values plotted in Figure 4E.
[c] Calculated from $\mu_{23}^{298K}$/298R and $h_{23}$/298R using Eq. 9. Propagated uncertainties calculated as previously described.[2]
[d] Predicted from the entropic contributions to the set of 4 two-way α-values at 25 °C (Table 2) and ASA information (Table S1). Propagated uncertainties calculated as previously described.[2]

**Table S3. Comparison of predicted $h_{23}$ (kcal mol$^{-1}$ molal$^{-1}$) from $\alpha^{h23}_{i,sp2C}$ with results of global fitting of $\Delta G^o_{tr}$ data**

| Solute | From global fitting of $\Delta G^o_{tr}$ vs. T to: | | Predicted from: |
|---|---|---|---|
| | Eq. 9 [a,b] | Eq. S4 [c] | Eq. 12 [b,d] |
| **urea** | -0.01 ± 0.06 | -0.12 ± 0.14 | -0.17 ± 0.41 |
| **methylurea** | 0.06 ± 0.14 | 0.17 ± 0.12 | 0.31 ± 0.31 |
| **ethylurea** | 0.07 ± 0.13 | 0.12 ± 0.13 | 0.50 ± 0.30 |
| **1,1-dmu** | 0.35 ± 0.1 | 0.46 ± 0.1 | 0.41 ± 0.28 |
| **1,3-dmu** | 0.51 ± 0.09 | 0.59 ± 0.09 | 0.79 ± 0.23 |
| **1,1-deu** | 1.45 ± 0.12 | 1.54 ± 0.12 | 0.83 ± 0.28 |
| **1,3-deu** | 1.55 ± 0.18 | 1.71 ± 0.18 | 1.17 ± 0.25 |
| **malonamide** | -0.14 ± 0.14 | -0.087 ± 0.14 | -0.43 ± 0.48 |
| **propionamide** | 0.11 ± 0.12 | 0.20 ± 0.12 | 0.34 ± 0.27 |
| **formamide** | -0.52 ± 0.09 | -0.49 ± 0.09 | -0.59 ± 0.39 |
| **methylFAD** | -0.36 ± 0.1 | -0.32 ± 0.1 | -0.12 ± 0.32 |
| **dimethylFAD** | 0.30 ± 0.14 | 0.40 ± 0.14 | 0.04 ± 0.30 |
| **acetamide** | -0.15 ± 0.14 | -0.15 ± 0.14 | -0.13 ± 0.31 |
| **methylacetamide** | -0.15 ± 0.15 | -0.03 ± 0.16 | 0.35 ± 0.25 |

[a] Obtained from $\mu_{23}^{298K}$ and 298 $s_{23}$ from Table 1 using Eq. 9.
[b] Values plotted in Figure 4D. Propagated uncertainties are calculated as previously described.[2]
[c] Obtained from $h_{23}$/298R from the fit to Eq. S4. Uncertainties are twice the fitting error.
[d] Predicted from enthalpic two-way α-values at 25 °C (Table 2) and ASA information (Table S1).

**Table S4. Comparison of $\mu_{233}$ values from two global fitting equations**

| Solute | $\mu_{233}$ (kcal mol$^{-1}$ molal$^{-2}$) [a] | |
|---|---|---|
| | From Eq. 8 [b] | From Eq. S4 [c] |
| **urea** | 0.000 ± 0.01 | 0.000 ± 0.01 |
| **methylurea** | 0.04 ± 0.01 | 0.04 ± 0.01 |
| **ethylurea** | 0.12 ± 0.01 | 0.11 ± 0.01 |
| **1,1-dmu** | 0.14 ± 0.02 | 0.15 ± 0.02 |
| **1,3-dmu** | 0.11 ± 0.01 | 0.11 ± 0.01 |
| **1,1-deu** | 0.14 ± 0.03 | 0.14 ± 0.03 |
| **1,3-deu** | 0.19 ± 0.03 | 0.19 ± 0.03 |
| **malonamide** | 0.10 ± 0.04 | 0.10 ± 0.03 |
| **propionamide** | 0.12 ± 0.03 | 0.12 ± 0.03 |
| **formamide** | 0.02 ± 0.01 | 0.02 ± 0.01 |
| **methylFAD** | 0.06 ± 0.01 | 0.06 ± 0.01 |
| **dimethylFAD** | 0.11 ± 0.01 | 0.11 ± 0.01 |
| **acetamide** | 0.02 ± 0.01 | 0.02 ± 0.01 |
| **methylacetamide** | 0.19 ± 0.02 | 0.19 ± 0.02 |

[a] Uncertainties reported as twice the fitting error.
[b] Obtained by fitting to Eq. 8 using Igor Pro 5.0.5A.
[c] Obtained by fitting to Eq. S4 using Igor Pro 5.0.5A.

**Table S5. Fitted vs. predicted amide-naphthalene $\mu_{23}$ values (kcal mol$^{-1}$ molal$^{-1}$) [a]**

| Solute | 25 °C | 25 °C | 10 °C | 10 °C | 45 °C | 45 °C |
|---|---|---|---|---|---|---|
| | Fitted (Eq.8) [b] | Predicted (Eq.10) [c] | Fitted (Eq.7) [d] | Predicted (Eq.10) [c] | Fitted (Eq.7) [d] | Predicted (Eq.10) [c] |
| **urea** | -0.15 ± 0.01 | - 0.14 ± 0.04 | -0.14 ± 0.01 | - 0.14 ± 0.05 | -0.16 ± 0.01 | - 0.14 ± 0.05 |
| **methylurea** | -0.37 ± 0.01 | - 0.37 ± 0.03 | -0.35 ± 0.01 | - 0.34 ± 0.04 | -0.40 ± 0.02 | - 0.42 ± 0.04 |
| **ethylurea** | -0.49 ± 0.01 | - 0.48 ± 0.03 | -0.46 ± 0.01 | - 0.43 ± 0.04 | -0.53 ± 0.02 | - 0.54 ± 0.04 |
| **1,1-dmu** | -0.61 ± 0.02 | - 0.55 ± 0.03 | -0.56 ± 0.02 | - 0.50 ± 0.03 | -0.68 ± 0.02 | - 0.62 ± 0.03 |
| **1,3-dmu** | -0.57 ± 0.01 | - 0.60 ± 0.03 | -0.52 ± 0.01 | - 0.53 ± 0.03 | -0.64 ± 0.01 | - 0.70 ± 0.03 |
| **1,1-deu** | -0.80 ± 0.02 | - 0.71 ± 0.03 | -0.68 ± 0.02 | - 0.64 ± 0.03 | -0.95 ± 0.02 | - 0.82 ± 0.03 |
| **1,3-deu** | -0.75 ± 0.02 | - 0.81 ± 0.03 | -0.63 ± 0.02 | - 0.71 ± 0.03 | -0.90 ± 0.02 | -0.95 ± 0.03 |
| **malonamide** | -0.36 ± 0.02 | - 0.34 ± 0.05 | -0.35 ± 0.02 | - 0.35 ± 0.05 | -0.38 ± 0.02 | -0.34 ± 0.06 |
| **propionamide** | -0.45 ± 0.02 | - 0.47 ± 0.03 | -0.42 ± 0.02 | - 0.43 ± 0.03 | -0.48 ± 0.03 | -0.53 ± 0.03 |
| **formamide** | -0.20 ± 0.01 | - 0.21 ± 0.04 | -0.21 ± 0.01 | - 0.23 ± 0.04 | -0.18 ± 0.01 | -0.18 ± 0.05 |
| **methylFAD** | -0.43 ± 0.01 | - 0.44 ± 0.03 | -0.43 ± 0.01 | - 0.43 ± 0.04 | -0.43 ± 0.01 | - 0.46 ± 0.04 |
| **dimethylFAD** | -0.67 ± 0.01 | - 0.64 ± 0.03 | -0.62 ± 0.02 | - 0.60 ± 0.04 | -0.73 ± 0.02 | - 0.68 ± 0.04 |
| **acetamide** | -0.31 ± 0.01 | - 0.40 ± 0.03 | -0.30 ± 0.01 | - 0.38 ± 0.04 | -0.32 ± 0.02 | - 0.42 ± 0.04 |
| **methylacetamide** | -0.63 ± 0.02 | - 0.63 ± 0.03 | -0.61 ± 0.02 | - 0.58 ± 0.03 | -0.66 ± 0.02 | - 0.69 ± 0.03 |

[a] Plotted in Figure 4A-C.
[b] Obtained by fitting to Eq. 8 with Igor Pro 5.0.5A. Uncertainties are twice the fitting error. Values at 25 °C reproduce those in Table 1.
[c] Predicted from the set of 4 two-way α-values (Table 2) and ASA information (Table S1) for amides and naphthalene. Propagated uncertainties are calculated as previously described.[2]
[d] Obtained by Eq. 7 from $\mu_{23}^{298K}$ and 298 $s_{23}$ from Table 1. Propagated uncertainties are calculated as previously described.[2]

**Table S6. Atom-atom amide-naphthalene interaction strengths $\alpha_{ij}^{\mu23}$ at 10, 25 and 45 °C and their entropic and enthalpic components (298 $\alpha_{ij}^{s23}$, $\alpha_{ij}^{h23}$)** [a]

| Atom-atom interaction | 10 °C $\alpha_{ij}^{\mu23}$ [b] | 25 °C $\alpha_{ij}^{\mu23}$ [b] | 45 °C $\alpha_{ij}^{\mu23}$ [b] | 298 $\alpha_{ij}^{s23}$ [c] | $\alpha_{ij}^{h23}$ [d] |
|---|---|---|---|---|---|
| **$sp^3C$-$sp^2C$** | - 8.8 ± 0.3 | - 10.6 ± 0.3 | - 12.9 ± 0.3 | 35 ± 2 | 24 ± 2 |
| **$sp^2O$-$sp^2C$** | - 16.7 ± 2.5 | - 11.9 ± 2.3 | - 5.4 ± 2.7 | - 96 ± 20 | - 110 ± 20 |
| **$sp^2N$-$sp^2C$** | 2.4 ± 0.9 | 0.7 ± 0.8 | - 1.5 ± 0.9 | 33 ± 7 | 34 ± 8 |
| **$sp^2C$-$sp^2C$** | - 3.8 ± 1.7 | - 5.3 ± 1.8 | - 7.3 ± 2.1 | 30 ± 15 | 25 ± 17 |
| **RMSE** [e] | 0.047 (in $\mu_{23}$) | 0.051 (in $\mu_{23}$) | 0.066 (in $\mu_{23}$) | 0.39 (in 298 $s_{23}$) | 0.37 (in $h_{23}$) |

[a] Atom-atom analysis using four $\alpha_{ij}^{\mu23}$ values. Units of $\alpha_{ij}^{\mu23}$, 298 $\alpha_{ij}^{s23}$, $\alpha_{ij}^{h23}$ are millical $mol^{-1}$ $molal^{-1}$ $Å^{-4}$. Propagated uncertainties are calculated as previously described.[2]
[b] Obtained from applying Eq. 10 to $\mu_{23}$ values from Table S5 at the given temperature, using ASA values from Table S1.
[c] Obtained from applying Eq. 11 to 298 $s_{23}$ values in the first column of Table S2, using ASA values from Table S1.
[d] Obtained from applying Eq. 12 to $h_{23}$ values in the first column of Table S3, using ASA values from Table S1.
[e] RMSE in $\mu_{23}$ calculated from Eq. S10. RMSE in $h_{23}$ and 298 $s_{23}$ calculated analogously. Units of RMSE are the same as $\mu_{23}$, $h_{23}$ and 298 $s_{23}$ (kcal $mol^{-1}$ $molal^{-1}$).

**Table S7. Contributions to $\mu_{23}$ , $h_{23}$ and 298 $s_{23}$ for interactions of amides with different C/O+N ratios, in kcal mol$^{-1}$ molal$^{-1}$. All values predicted from two-way alpha values (Table 2) and ASA information (Table S1).**

| Amide | $\mu_{23}$ [a] | $sp^3C$-$sp^2C$ | $sp^2O$-$sp^2C$ | $sp^2N$-$sp^2C$ | $sp^2C$-$sp^2C$ |
|---|---|---|---|---|---|
| urea | -0.14 ± 0.06 | No $sp^3C$ | -0.16 ± 0.03 | 0.03 ± 0.03 | -0.01 ± 0.01 |
| formamide | -0.21 ± 0.07 | No $sp^3C$ | -0.17 ± 0.03 | 0.01 ± 0.02 | -0.06 ± 0.02 |
| malonamide | -0.34 ± 0.08 | -0.14 ± 0.01 | -0.21 ± 0.04 | 0.02 ± 0.03 | -0.01 ± 0.01 |
| 1,1-deu | -0.71 ± 0.05 | -0.6 ± 0.02 | -0.12 ± 0.02 | 0.01 ± 0.01 | -0.01 ± 0.01 |
| 1,3-deu | -0.81 ± 0.05 | -0.72 ± 0.02 | -0.09 ± 0.02 | 0.01 ± 0.01 | -0.01 ± 0.01 |
| **Amide** | **298 $s_{23}$ [b]** | **$sp^3C$-$sp^2C$** | **$sp^2O$-$sp^2C$** | **$sp^2N$-$sp^2C$** | **$sp^2C$-$sp^2C$** |
| urea | -0.03 ± 0.54 | No $sp^3C$ | -1.26 ± 0.26 | 1.17 ± 0.26 | 0.06 ± 0.03 |
| formamide | -0.38 ± 0.58 | No $sp^3C$ | -1.35 ± 0.28 | 0.64 ± 0.14 | 0.33 ± 0.16 |
| malonamide | -0.08 ± 0.66 | 0.46 ± 0.03 | -1.73 ± 0.35 | 1.11 ± 0.24 | 0.07 ± 0.03 |
| 1,1-deu | 1.39 ± 0.37 | 1.69 ± 0.1 | -0.75 ± 0.15 | 0.4 ± 0.09 | 0.05 ± 0.02 |
| 1,3-deu | 1.54 ± 0.43 | 2.00 ± 0.12 | -0.94 ± 0.19 | 0.45 ± 0.1 | 0.03 ± 0.02 |
| **Amide** | **$h_{23}$ [c]** | **$sp^3C$-$sp^2C$** | **$sp^2O$-$sp^2C$** | **$sp^2N$-$sp^2C$** | **$sp^2C$-$sp^2C$** |
| urea | -0.17 ± 0.61 | No $sp^3C$ | -1.41 ± 0.29 | 1.19 ± 0.28 | 0.05 ± 0.03 |
| formamide | -0.59 ± 0.65 | No $sp^3C$ | -1.51 ± 0.31 | 0.65 ± 0.15 | 0.27 ± 0.18 |
| malonamide | -0.43 ± 0.73 | 0.32 ± 0.03 | -1.94 ± 0.4 | 1.13 ± 0.27 | 0.06 ± 0.04 |
| 1,1-deu | 0.83 ± 0.48 | 1.4 ± 0.14 | -1.05 ± 0.22 | 0.46 ± 0.11 | 0.02 ± 0.02 |
| 1,3-deu | 1.17 ± 0.43 | 1.67 ± 0.16 | -0.85 ± 0.17 | 0.31 ± 0.07 | 0.04 ± 0.03 |

[a] Predicted value (Table S5).
[b] Predicted value (Table S2).
[c] Predicted value (Table S3).

**Table S8. CH–π and π-π amide-naphthalene interaction strengths $\alpha_{ij}^{\mu 23}$ at 10, 25 and 45 °C and their entropic and enthalpic components (298 $\alpha_{ij}^{s23}$, $\alpha_{ij}^{h23}$) [a]**

| Interaction | 10 °C $\alpha_{ij}^{\mu 23}$ [b,c] | 25 °C $\alpha_{ij}^{\mu 23}$ [b,c] | 45 °C $\alpha_{ij}^{\mu 23}$ [b,c] | 298 $\alpha_{ij}^{s23}$ [b,d] | $\alpha_{ij}^{h23}$ [b,e] |
|---|---|---|---|---|---|
| **CH-π** | - 9.9 ± 0.2 | - 11.3 ± 0.2 | - 13.2 ± 0.2 | 27.6 ± 1.3 | 16.3 ± 1.4 |
| **π-π** | - 3.9 ± 0.2 | - 3.6 ± 0.2 | - 3.2 ± 0.2 | - 5.5 ± 1.4 | - 9.1 ± 1.4 |
| **RMSE** [f] | 0.066 (in $\mu_{23}$) | 0.061 (in $\mu_{23}$) | 0.066 (in $\mu_{23}$) | 0.42 (in 298 $s_{23}$) | 0.43 (in $h_{23}$) |

[a] Analysis using two $\alpha_{ij}$ values for CH–π and π-π interactions (Eq. S5, S7, S8) for comparison with atom-atom results in Table S6 for amide-naphthalene interactions.
[b] Units of $\alpha_{ij}^{\mu 23}$, 298 $\alpha_{ij}^{s23}$ and $\alpha_{ij}^{h23}$ are millical $mol^{-1}$ $molal^{-1}$ $Å^{-4}$. Propagated uncertainties are calculated as previously described.[2] See Figure S7.
[c] Obtained from applying Eq. S5 to $\mu_{23}$ values from Table S5 at the given temperature, using ASA values from Table S1.
[d] Obtained from applying Eq. S7 to 298 $s_{23}$ values in the first column of Table S2 , using ASA values from Table S1.
[e] Obtained from applying Eq. S8 to $h_{23}$ values in the first column of Table S3, using ASA values from Table S1.
[f] RMSE in $\mu_{23}$ calculated from Eq. S10. RMSE in $h_{23}$ and 298 $s_{23}$ calculated analogously. Units of RMSE are the same as $\mu_{23}$, $h_{23}$ and 298 $s_{23}$ (kcal $mol^{-1}$ $molal^{-1}$).

**Table S9. Comparison of two-way $\alpha_{ij}$ values for different fits of amide-aromatic and/or amide-amide $\mu_{23}$ values (all $\alpha_{ij}$ values in millical mol$^{-1}$ molal$^{-1}$ Å$^{-4}$)**

| | 14 [a] (or 24 [a,b]) amide-aro $\mu_{23}$ 4 $\alpha_{ij}$ [c] | 14 [a] (or 24 [a,b]) amide-aro $\mu_{23}$ 2 $\alpha_{ij}$ [d] | 109 $\mu_{23}$ [a,b,e] 10 $\alpha_{ij}$ [e] | 109 $\mu_{23}$ [a,b,e] 4 $\alpha_{ij}$ [f] |
|---|---|---|---|---|
| **sp$^3$C-sp$^3$C** | | | - 4.0 ± 0.1 | - 1.7 ± 0.1 |
| **sp$^3$C-aro π** | | - 11.3 ± 0.2 (- 11.6 ± 0.2) | | - 11.8 ± 0.1 |
| **sp$^3$C-sp$^2$C** | - 10.6 ± 0.3 (- 10.8 ± 1.6) | | - 10.8 ± 0.4 | |
| **sp$^3$C-am π** | | | | 0.0 ± 0.2 |
| **sp$^3$C-sp$^2$O** | | | 11.5 ± 0.5 | |
| **sp$^3$C-sp$^2$N** | | | - 4.0 ± 0.2 | |
| **π – π** | | | | - 2.8 ± 0.1 |
| **am π- am π** | | | | |
| **aro π- am π** | | - 3.6 ± 0.2 (- 3.2 ± 0.3) | | |
| **sp$^2$C-sp$^2$O** | - 11.9 ± 2.3 (- 11.4 ± 3.2) | | - 11.4 ± 4.3 | |
| **sp$^2$C-sp$^2$N** | 0.7 ± 0.8 (0.5 ± 0.4) | | 0.5 ± 1.6 | |
| **sp$^2$C-sp$^2$C** | - 5.3 ± 1.8 (- 5.1 ± 4.4) | | - 5.0 ± 3.1 | |
| **RMSE in $\mu_{23}$ [g]** | 0.051 (0.048) | 0.061 (0.056) | 0.024 | 0.033 |

[a] This work
[b] SI Ref. 3
[c] Eq. 10
[d] Eq. S5
[e] SI Ref. 7
[f] Eq. S9
[g] Eq. S10. Units of RMSE are the same as $\mu_{23}$ (kcal mol$^{-1}$ molal$^{-1}$).

**Table S10. Enthalpy-entropy relationships observed in studies of solute effects on protein unfolding thermodynamics**

| At 25 °C | **Concentration scale** | **$\Delta\mu_{23}$ (kcal mol$^{-1}$ per unit concentration)** | **$\Delta h_{23}$ (kcal mol$^{-1}$ per unit concentration)** | **$298\Delta s_{23}$ (kcal mol$^{-1}$ per unit concentration) [a]** |
|---|---|---|---|---|
| **Urea – HPr unfolding [b]** | Molar | - 1.13 ± 0.04 | - 2.9 ± 1.3 | (- 1.8 ± 1.3 ) |
| **Urea -lac DBD unfolding [c]** | Molar | - 0.45 ± 0.01 | - 0.66 ± 0.09 | - 0.20 ± 0.09 |
| **GB – lac DBD unfolding [d]** | Molar | 0.75 ± 0.05 | 0 ± 0.2 | - 0.75 ± 0.33 |
| **EG – SH3 unfolding [e]** | Molar | 0.28 ± 0.05 | 4.1 ± 0.1 | (3.8 ± 0.1) |
| **EG – SH3 unfolding [e]** | Weight (g/mL) | 4.5 ± 0.8 | 66 ± 11 | (62 ± 12) |
| **PEG400 – SH3 unfolding [e]** | Weight (g/mL) | 2.1 ± 0.4 | 60 ± 4 | (58 ± 5) |

[a] Values in parentheses calculated from $298\ \Delta s_{23} = \Delta h_{23} - \Delta\mu_{23}$

[b] SI Ref. 10

[c] GB: Glycine betaine. See SI Ref. 11

[d] SI Ref. 12

[e] EG, Ethylene glycol; PEG 400, ethylene glycol oligomers with an average molecular weight of 400 g/mol. Tabulated information obtained from Fig 1 of SI Ref. 13.

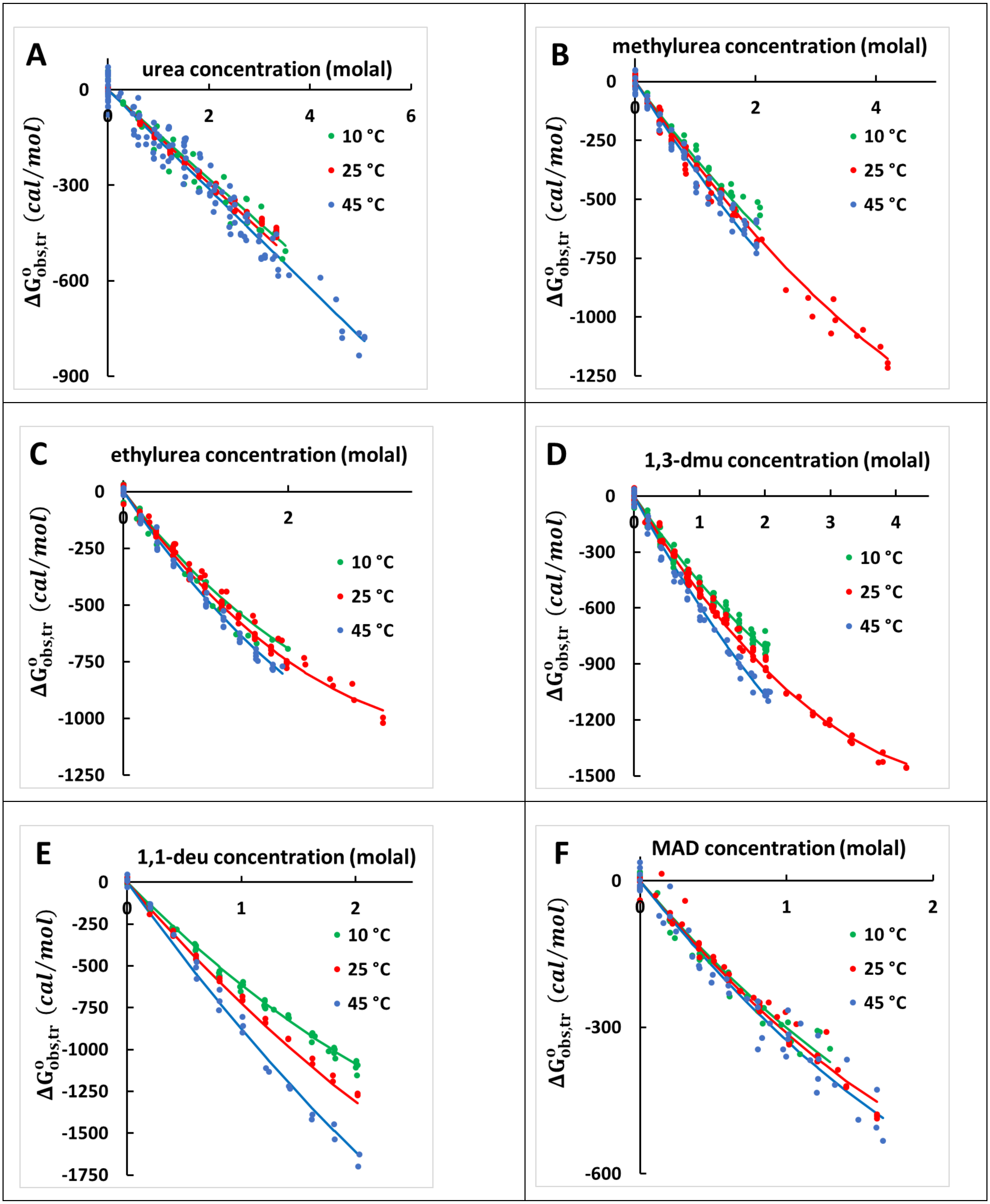

A
urea concentration (molal)
0
2
4
6
-300
-600
-900
$\Delta G^o_{obs,tr}$ (cal/mol)
10 °C
25 °C
45 °C
B
methylurea concentration (molal)
0
2
4
-250
-500
-750
-1000
-1250
$\Delta G^o_{obs,tr}$ (cal/mol)
10 °C
25 °C
45 °C
C
ethylurea concentration (molal)
0
2
-250
-500
-750
-1000
-1250
$\Delta G^o_{obs,tr}$ (cal/mol)
10 °C
25 °C
45 °C
D
1,3-dmu concentration (molal)
0
1
2
3
4
-300
-600
-900
-1200
-1500
$\Delta G^o_{obs,tr}$ (cal/mol)
10 °C
25 °C
45 °C
E
1,1-deu concentration (molal)
0
1
2
-250
-500
-750
-1000
-1250
-1500
-1750
$\Delta G^o_{obs,tr}$ (cal/mol)
10 °C
25 °C
45 °C
F
MAD concentration (molal)
0
1
2
-300
-600
$\Delta G^o_{obs,tr}$ (cal/mol)
10 °C
25 °C
45 °C

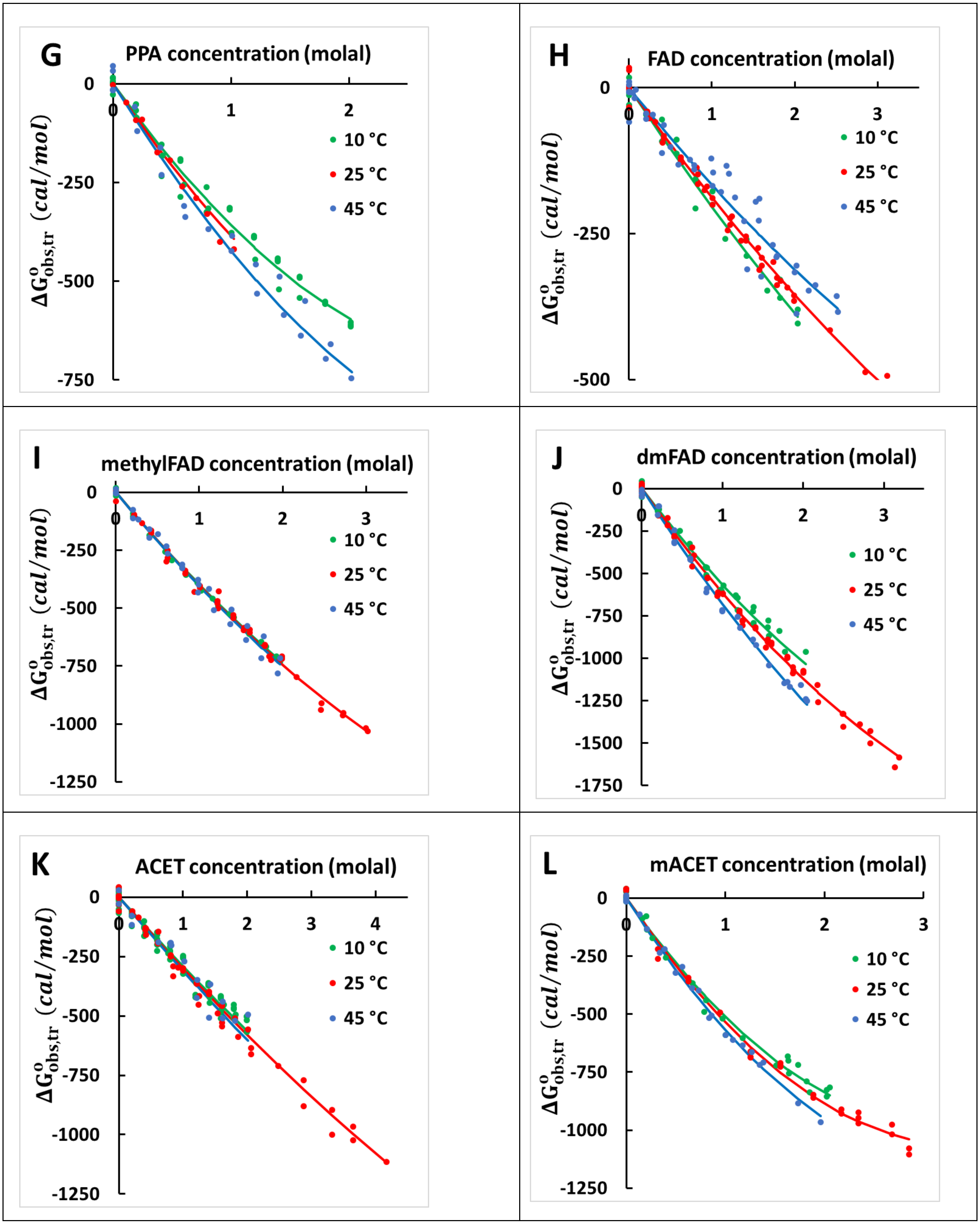
G
PPA concentration (molal)
0
-250
-500
-750
0
1
2
10 °C
25 °C
45 °C
$\Delta G^{o}_{obs,tr}$ $(cal/mol)$
H
FAD concentration (molal)
0
-250
-500
0
1
2
3
10 °C
25 °C
45 °C
$\Delta G^{o}_{obs,tr}$ $(cal/mol)$
I
methylFAD concentration (molal)
0
-250
-500
-750
-1000
-1250
0
1
2
3
10 °C
25 °C
45 °C
$\Delta G^{o}_{obs,tr}$ $(cal/mol)$
J
dmFAD concentration (molal)
0
-250
-500
-750
-1000
-1250
-1500
-1750
0
1
2
3
10 °C
25 °C
45 °C
$\Delta G^{o}_{obs,tr}$ $(cal/mol)$
K
ACET concentration (molal)
0
-250
-500
-750
-1000
-1250
0
1
2
3
4
10 °C
25 °C
45 °C
$\Delta G^{o}_{obs,tr}$ $(cal/mol)$
L
mACET concentration (molal)
0
-250
-500
-750
-1000
-1250
0
1
2
3
10 °C
25 °C
45 °C
$\Delta G^{o}_{obs,tr}$ $(cal/mol)$

**Figure S1. Analysis of effects of amide compounds on naphthalene solubility at 10, 25 and 45 °C.** Values of $\Delta G^{o}_{obs,tr}$, the observed standard free energy of transfer of naphthalene from water to the amide solution at concentration $m_3$ ($\Delta G^{o}_{obs,tr} = -RT\ln\left(m_2^{ss,m3}/m_2^{ss,0}\right)$; Eq. 1), are plotted as a function of amide concentration $m_3$ at 10 °C (green), 25 °C (red), and 45 °C (blue). In each panel, plotted curves are from a global fit to data at 10, 25 and 45 °C (Eq. 8; fitting parameters listed in SI Tables S4-5; see also Figure 1). Abbreviations: dmu, dimethylurea; deu, diethylurea; MAD, malonamide; PPA, propionamide; FAD, formamide; dmFAD, dimethylformamide; ACET, acetamide; mACET, methylacetamide.

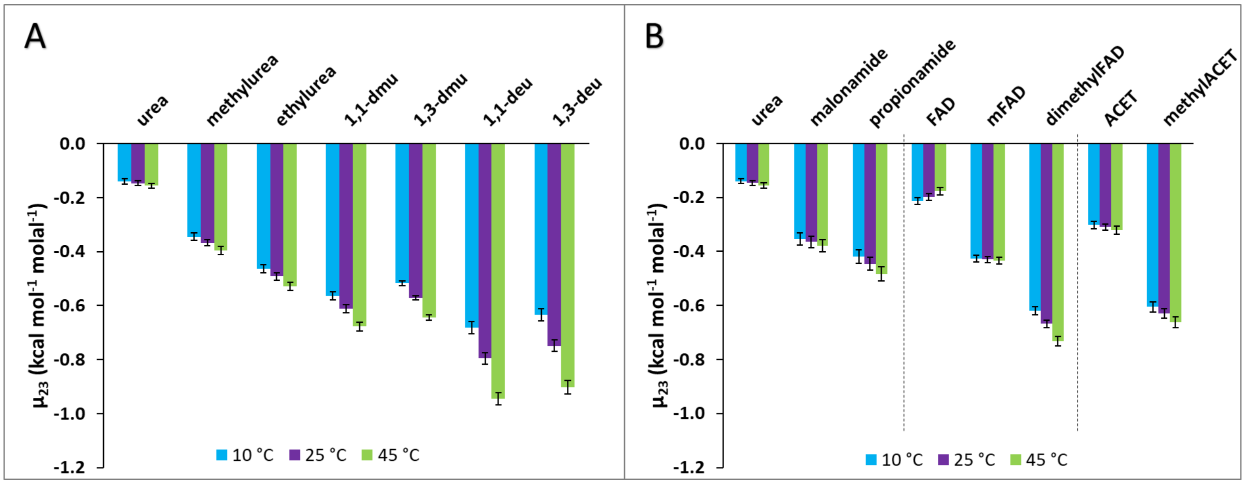


**Figure S2. Bar graph summary of preferential interactions ($\mu_{23}$) of amide compounds with naphthalene at 10, 25, and 45 °C.** Values of $\mu_{23}$ at 25 °C are from the global analysis (Eq. 8; Table 1) assuming $s_{23}$ and $\mu_{233}$ are independent of temperature. The $\mu_{23}$-values at 10 and 45 °C are calculated from Eq. 7 and listed in Table S5. Panel A shows the series of alkylated ureas in order of increasing C/(O+N) ASA ratio. Panel B shows three smaller groups of amides, each also ranked in order of increasing C/(O+N) ASA ratio.

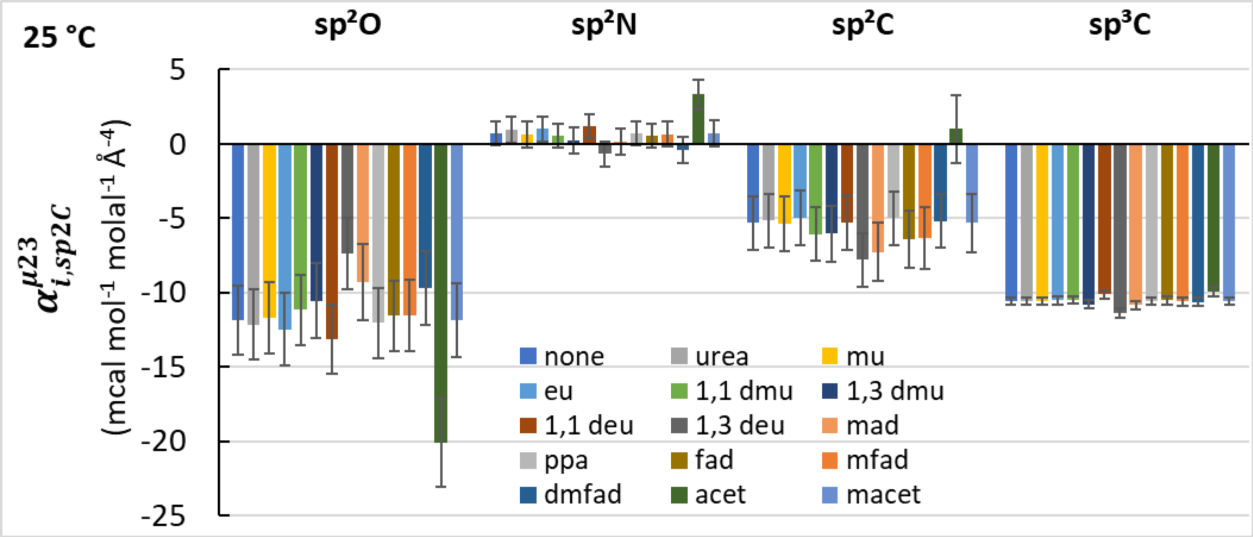


**Figure S3. Effect on $\alpha^{\mu23}_{i,sp2C}$ for interactions of amide atoms with aromatic $sp^2C$ atoms at 25 °C when one amide compound is removed from the 14 amide data set.** Values of $\alpha^{\mu23}_{i,sp2C}$ are shown with propagated uncertainties, calculated as previously described.[2] The index i stands for either amide $sp^2O$, amide $sp^2N$, amide $sp^2C$ or aliphatic $sp^3C$ atoms.

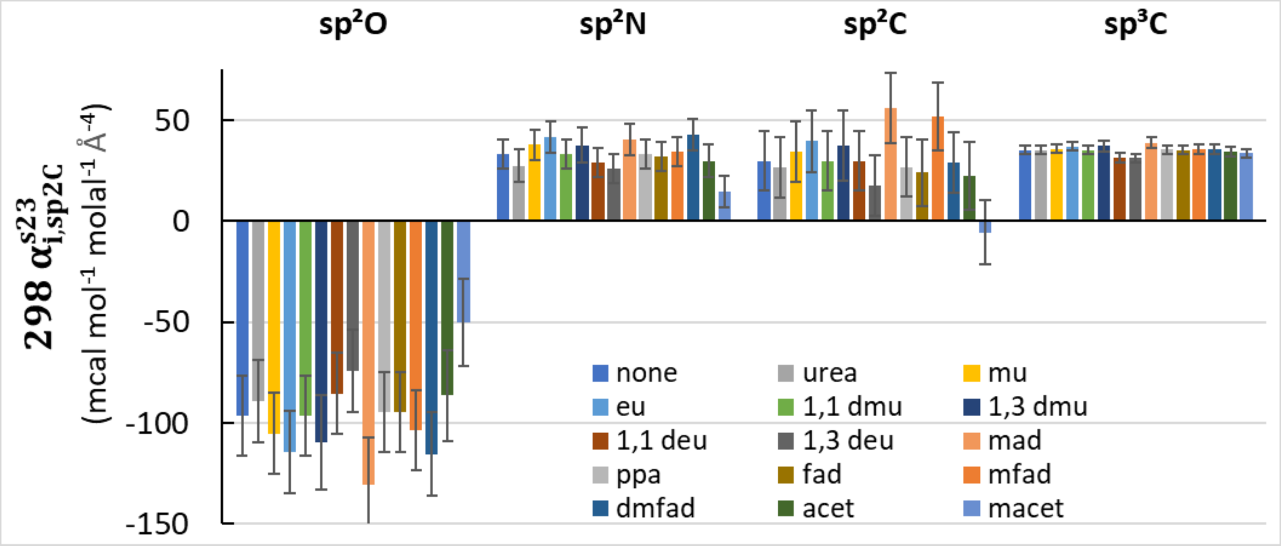


**Figure S4. Effect on 298 $\alpha^{s23}_{i,sp2C}$ for interactions of amide atoms with aromatic $sp^2C$ atoms at 25 °C when one amide compound is removed from the 14 amide data set.** Values of 298 $\alpha^{s23}_{i,sp2C}$ are shown with propagated uncertainties, calculated as previously described.[2] The index i stands for either amide $sp^2O$, amide $sp^2N$, amide $sp^2C$ or aliphatic $sp^3C$ atoms.

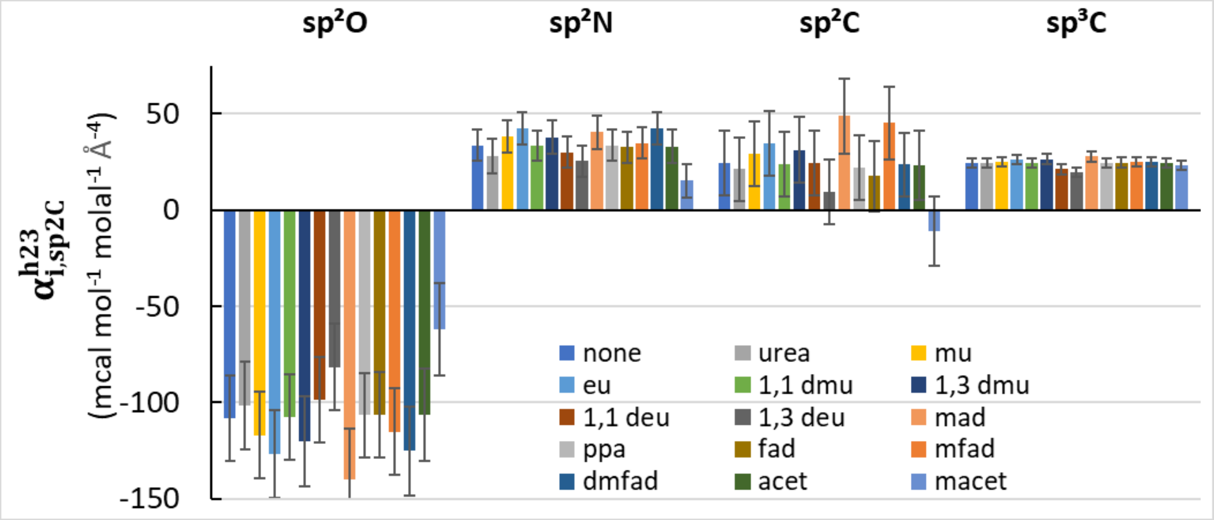


**Figure S5. Effect on $\alpha^{h23}_{i,sp2C}$ for interactions of amide atoms with aromatic $sp^2C$ atoms at 25 °C when one amide compound is removed from the 14 amide data set.** Values of $\alpha^{h23}_{i,sp2C}$ are shown with propagated uncertainties, calculated as previously described.[2] The index i stands for either amide $sp^2O$, amide $sp^2N$, amide $sp^2C$ or aliphatic $sp^3C$ atoms.

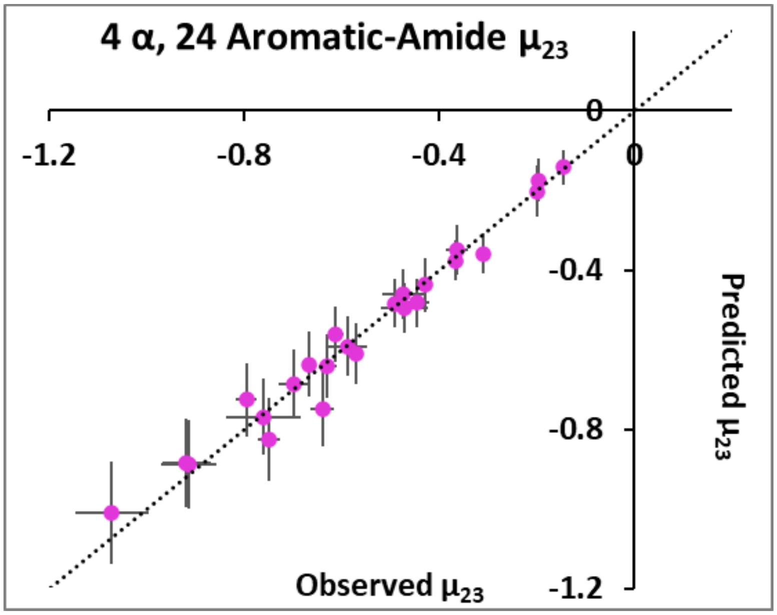


**Figure S6. Comparison of observed $\mu_{23}$ values quantifying 24 amide-aromatic interactions with predictions from atom-atom (4 $\alpha_{ij}$-value) analysis.** Predictions use Eq. 10 and ASA information from Table S1. Values of $\alpha_{ij}$ used in these predictions are from Table S9. In all cases units of $\mu_{23}$ are kcal mol$^{-1}$ molal$^{-1}$.

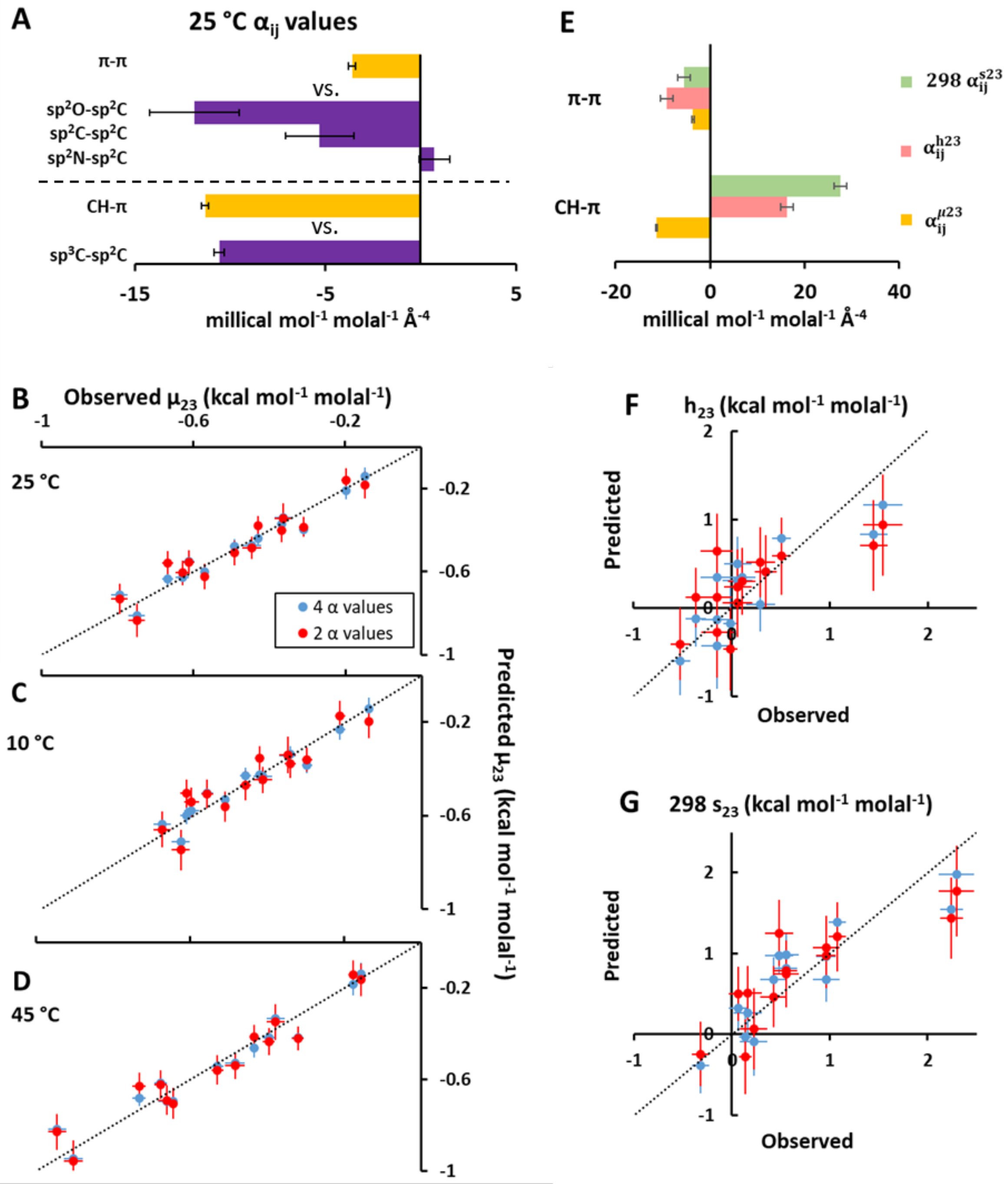


**Figure S7. Analysis of amide-naphthalene interactions as π-π/CH-π or amide atom-naphthalene atom interactions.** Panel A) Free energy π-π and CH-π alpha values, for comparison with the corresponding atom-atom information in Figure 3B. Panels B-D) Comparisons of observed 25, 10 and 45 °C $\mu_{23}$ values with those predicted from π-π/CH-π analysis (red: 2 $\alpha$-values, Eq. S5-6) and atom-atom analysis (blue: 4 $\alpha$-values, Eq.10). Panel E) Entropic and enthalpic components of the free energy π-π and CH-π $\alpha$-values (Eqs. S7, S8), for comparison with the corresponding atom-atom information in Figure 3B. Panels F-G) Comparisons of observed $h_{23}$ and 298 $s_{23}$ values with those predicted from π-π/CH-π analysis (red: 2 $\alpha$-values, Eqs. S7, S8) and atom-atom analysis (blue: 4 $\alpha$-values, Eqs. 11-12).

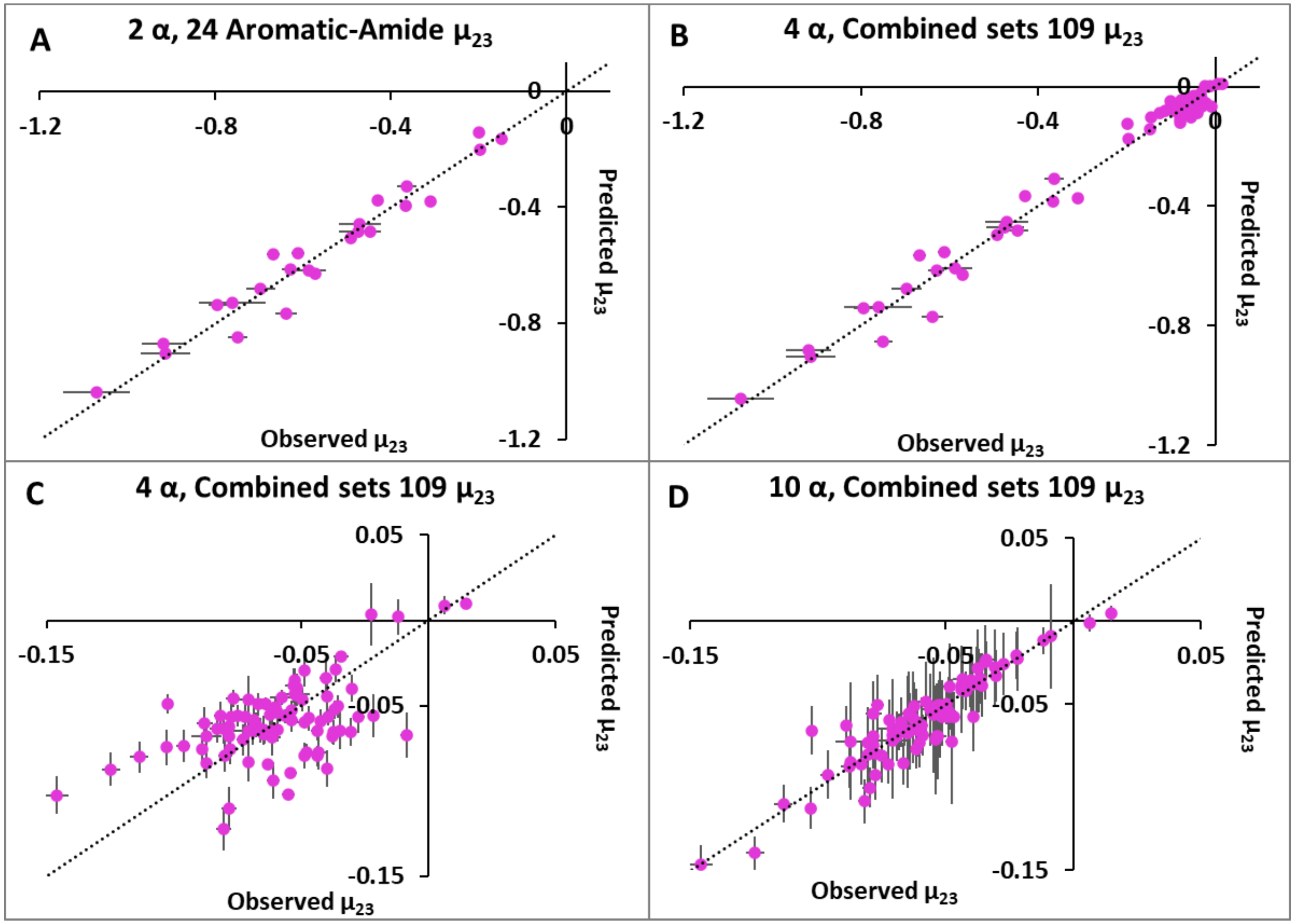


**Figure S8. Predicted vs. observed 25 °C $\mu_{23}$ values (in kcal mol$^{-1}$ molal$^{-1}$) for analyses based on π-π and CH-π interactions instead of atom-atom interactions.** Panel A) $\mu_{23}$ values for 24 amide-aromatic interactions, predicted using 2 $\alpha_{ij}$-values (Eqs. S5-6; see Table S9). Panel B) $\mu_{23}$ values for combined set of 85 amide-amide and 24 amide-aromatic interactions (109 $\mu_{23}$ values), predicted using 4 $\alpha_{ij}$-values (Eq. S9; see Table S9). Panel C) Expanded scale plot of $\mu_{23}$ values for amide-amide portion of Panel B. Panel D) An expanded scale plot of $\mu_{23}$ values for amide-amide portion of Figure 5B using the atom-atom (10 $\alpha_{ij}$-value) fit, for comparison with the 4 $\alpha_{ij}$-value fit in Panel C. ASA information for all panels is from Table S1. RMSE values (Eq. S10) in Table S9 provide a statistical comparison of the quality of these predictions.
